# De novo design of diverse small molecule binders and sensors using Shape Complementary Pseudocycles

**DOI:** 10.1101/2023.12.20.572602

**Authors:** Linna An, Meerit Said, Long Tran, Sagardip Majumder, Inna Goreshnik, Gyu Rie Lee, David Juergens, Justas Dauparas, Ivan Anishchenko, Brian Coventry, Asim K. Bera, Alex Kang, Paul M. Levine, Valentina Alvarez, Arvind Pillai, Christoffer Norn, David Feldman, Dmitri Zorine, Derrick R. Hicks, Xinting Li, Mariana Garcia Sanchez, Dionne K. Vafeados, Patrick J. Salveson, Anastassia A. Vorobieva, David Baker

## Abstract

A general method for designing proteins to bind and sense any small molecule of interest would be widely useful. Due to the small number of atoms to interact with, binding to small molecules with high affinity requires highly shape complementary pockets, and transducing binding events into signals is challenging. Here we describe an integrated deep learning and energy based approach for designing high shape complementarity binders to small molecules that are poised for downstream sensing applications. We employ deep learning generated psuedocycles with repeating structural units surrounding central pockets; depending on the geometry of the structural unit and repeat number, these pockets span wide ranges of sizes and shapes. For a small molecule target of interest, we extensively sample high shape complementarity pseudocycles to generate large numbers of customized potential binding pockets; the ligand binding poses and the interacting interfaces are then optimized for high affinity binding. We computationally design binders to four diverse molecules, including for the first time polar flexible molecules such as methotrexate and thyroxine, which are expressed at high levels and have nanomolar affinities straight out of the computer. Co-crystal structures are nearly identical to the design models. Taking advantage of the modular repeating structure of pseudocycles and central location of the binding pockets, we constructed low noise nanopore sensors and chemically induced dimerization systems by splitting the binders into domains which assemble into the original pseudocycle pocket upon target molecule addition.

**One Sentence Summary:** We use a pseuodocycle-based shape complementarity optimizing approach to design nanomolar binders to diverse ligands, including the flexible and polar methotrexate and thyroxine, that can be directly converted into ligand-gated nanopores and chemically induced dimerization systems.

## Main Text

The design of small molecule (SM) binding proteins is more challenging than the design of protein binders as there are fewer atoms to interact with; hence high shape complementary (SC) pockets which make contact with the majority of the available atoms are required for high affinity(*1*). Designing such complementary pockets requires high accuracy modeling of protein-ligand packing, hydrogen bonding, π-π and other interactions, and the generation of scaffolds that can harbor such close interactions surrounding a large fraction of the SM surface. These challenges are particularly critical for designing binders to flexible polar compounds: polar groups make hydrogen bonds to water in the unbound state, which must be replaced by hydrogen bonds to the protein for binding to be favorable, and flexible compounds lose considerable entropy upon binding which must be compensated by extensive favorable interactions. In nature, SM binding receptors not only specifically bind their targets but transduce binding events into downstream signals(*2*); a further design challenge is to devise general approaches for similarly coupling binding to sensing. Previous successes in SM binder design have focused on relatively rigid hydrophobic targets and often have required considerable experimental optimization to improve binding affinities from micromolar level to high nanomolar levels(*3*–*8*). Deep learning (DL) diffusion approaches have been extended to SM binder design, but successes thus far are all for large, bulky, rigid nonpolar molecules(*9*); the power of DL approaches may be limited by the significantly smaller amount of training data compared to the protein-protein interaction case(*9*, *10*). Overall, generation of binders to polar and flexible small molecules, and systematic approaches for converting such binders into sensors, remain largely outstanding challenges for computational protein design.

We set out to develop a general method for designing SM binding proteins with high shape complementarity for their targets and with properties enabling facile conversion into sensors. We hypothesized that a design approach beginning with identification of protein scaffolds with high SC(*1*, *11*) to the targeted SM would be able to ultimately achieve higher affinity binding than approaches based on fixed scaffolds(*3*, *5*, *6*, *8*), and enable binding to flexible and polar targets. We reasoned further that if the scaffolds could be split into two or more independently folded domains such that SM binding energy drives association, binders could be readily converted into sensors. This strategy requires that the domains be folded prior to association to avoid aggregation and proteolysis of the split domains in the unbound state. Thus scaffold sets that can harbor binding pockets of widely ranging shapes and sizes, and at the same time can be readily split into multiple independently folded domains could provide a general solution to the sensor design problem(*11*).

We reasoned that designed pseudocyclic scaffolds consisting of a repeating structural unit surrounding a central pore or pocket could satisfy the above requirements(*11*) (**Fig 1a**). First, depending on the geometry of the structural unit and the number of units in the closed pseudocycle, the central pocket can have a very wide range of sizes and shapes. Second, since the interactions within the protein are almost entirely local–between residues close along the sequence within single repeat units or between adjacent repeat units, pseudocycles can be split into multiple domains that retain almost all of the stabilizing interactions present in the unsplit protein, and hence these split domains are likely to fold and be well behaved on their own(*11*). We set out to (**Fig 1a** & **S1a-b**) develop a general approach integrating DL and Rosetta(*12*) energy-based design methods(*11*, *13*–*16*) to generate high SC pseudocycle based binding proteins for any desired small molecule (**Fig 1b**).

**Figure 1.**
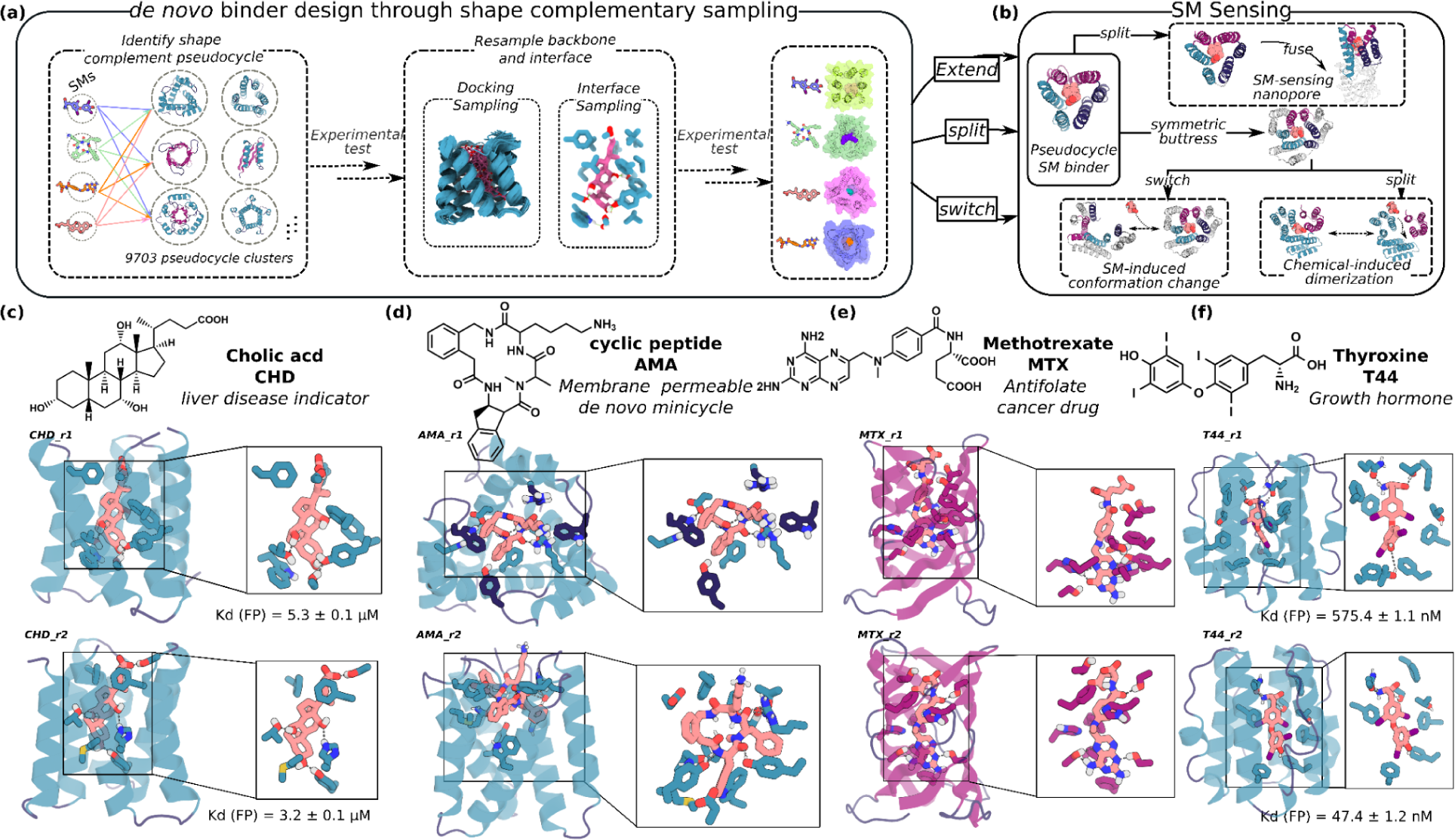
Pseudocycle-based SC optimizing design method and target SMs. **(a)** Diverse conformers of the SM of interest are docked into deep learning generated pseudocycles containing a wide array of central pockets(*11*), and the interface sequence optimized for high affinity binding using Rosetta or LgandMPNN. Top ranked designs are tested experimentally, and the backbones of the best hits and the docked poses are extensively resampled. Following sequence design, top ranked second round designs are experimentally tested (see **Fig S1**). **(b)** Because pseudocycles are constructed from modular repeating units which surround the central binding pocket, the binders can be readily transformed into sensors through multiple strategies. **(c-f)** Examples of first round design models for each target ligand.

We chose four SMs as binding targets: cholic acid (CHD), methotrexate (MTX), thyroxine (T44), and a de novo designed cell permeable cyclic tetrapeptide called AMA (**Fig. 1, S4 a-d,** & **Table S4**). CHD is the primary bile acid, and detection of its free form is important for liver disease determination(*17*). MTX is an anti-folate cancer treatment agent which requires regular blood monitoring to reduce adverse outcomes for patients(*18*). T44 is a critical human hormone regulating energy usage and other functions; at home monitoring to detect free T44 levels is used for patient thyroid condition diagnosis. The current commercially available detection methods for CHD, MTX, and T44 are either time-consuming, chromatography-based analytical methods, or immunoassays(*17*–*21*), which cannot distinguish individual molecules from variants (CHD, T44) or bound from free forms (T44, MTX); higher affinity and specificity binders are needed for rapid at-home SM sensing devices. From the design perspective, T44 and MTX are significantly more flexible and polar than previous SM targets for computational protein design(*3*–*8*), and MTX is particularly challenging due to its high polarity and flexibility (**Fig S4e** & **Table S4**). To test the ability of the method to design binders to larger ligands, we also included AMA (**Fig 1** & **S4a**), a de novo designed cell membrane-permeable tetrapeptidic macrocycle(*22*); designed binders for such compounds could be turned into chemical-induced dimerization (CID) systems enabling bioorthogonal control for adoptive cell therapies and other applications. We obtained or synthesized fluorescein (FITC, see ‘***Preparation of the FITC-labeled SM target***’ in Methods) or biotin labeled versions of all four ligands for experimental screening (**Fig S4**).

### Identification of pseudocycle scaffolds with high shape complementary to target

We begin by identifying for a given ligand of interest the most shape complementary pockets present in AlphaFold2 and ProteinMPNN generated pseudocycle scaffolds(*11*) with a wide range of pocket shapes and sizes (**Fig 1a** & **S1a**). We first used RDKit(*23*) to generate hundreds of physically allowed rotamers for each ligand, clustered them based on all-atom root-mean-square deviation (r.m.s.d), and selected a low Rosetta energy conformer to represent each cluster (it is not sufficient to only consider the lowest energy conformer as this overly limits the maximal shape complementarity that can be achieved; many native SM binders bind the ligand in a conformation different from its free form). For each ligand, we docked one to a few hundred rotamers (**Table S1**) to the pockets in 9,703 pseudocycles(*11*) using Rifgen/Rifdock(*3*), generating in total 1-10 million docks. To rapidly identify docks with relatively high SC between the protein and SM, we developed an *in silico* two-step ‘predictor’ (**Fig S1a**) which benchmarks showed highly correlated with the SC after the much more expensive full sequence design (**Fig S5**). The ‘predictor’ first uses a quick Rosetta protocol (∼10 dock/CPU second, see ‘***Predictor to select best docks****’* in Methods) to pack the ligand-protein interface and estimate the protein-ligand contacts (using ‘contact_molarcular_surface’, CMS); The best solutions found in this screen were subjected to a more thorough interface design and scoring protocol (∼1 dock/CPU second), and additional features including the predicted binding energy and the number of hydrogen bonds to the ligand were used to select promising docks (see **‘*Step-wise SM binder design pipeline in detail*’** in Methods). Full fixed-backbone sequence design was carried out for the top selected ∼100,000 docks (**Table S1**) using position specific scoring matrix (PSSM)-based Rosetta design(*12*) (see “***PSSM-based Rosetta design protocol****”* in Methods) or using a new deep learning based ligandMPNN sequence design method which enables direct optimization of interactions with small molecules (see “***Iterative ligandMPNN design protocol****”* in Methods)(*16*) (**Table S2**). 5-10,000 designs for each target predicted to fold to the designed structures by AlphaFold2(*14*) (AF2) and make low free energy interactions with the ligand (as computed by Rosetta) were selected for experimental characterization (**Table S1**).

The selected designs were encoded in oligonucleotide libraries, displayed on the yeast cell surface and binding was assessed using fluorescence-activated cell sorting (FACS), and deep sequencing (see ***‘High-throughput methods for binder identification’*** in Methods). For CHD (**Fig 1c** & **S2a-b**) we obtained two binders based from the same scaffold with Kds of 3.2 ± 0.1 and 5.3 ± 0.1 μM by fluorescence polarization (FP, see ‘Fluorescence polarization studies’ in Methods); in the design models all ligand polar atoms are contacted by sidechains, with these key polar residues buttressed by hydrogen-bonding networks (**Fig 1c** & **S2a-b**). For T44, we again obtained two binders based on the same scaffold with Kds of 575.4 ± 1.1 nM and 47.4 ± 2 nM using FP (**Fig 1f** & **S2c-d**). For MTX and AMA, we identified one and two potential binders from each two and four hit pseudocycle scaffolds, respectively (**Fig 1d-e, S3** & **S6**); these binders were weaker than the CHD and T44 binders but still showed clear ligand-specific binding signals on FACS using yeast surface display (**Fig S3**). In all four cases, there were 100 or fewer designs from the scaffolds that gave rise to binders (for example, only 32 designs were tested from the scaffold for CHD which yielded 2 binders), so had we been able to identify the most appropriate scaffold for each compound in advance, the protocol would have yielded binders with small scale screening (the contact molecular surface with ligand was generally higher for the designs with binding activity, but more data is needed to set thresholds; **Fig S1c**). Binders were obtained using both the PSSM-based Rosetta design protocol and the ligandMPNN design protocol (**Table S2**), suggesting that when there is close shape complementarity between protein and ligand, the details of the sequence design method become less critical.

We were able to obtain a co-crystal structure of one of the CHD binders in complex with CHD (CHD_r1, **Fig 2** & **S7a-c**). The co-crystal structure agrees closely with the computational design model with a 0.80 Å Cɑ-r.m.s.d between the two. All ligand-protein hydrogen bond interactions and hydrogen bond networks were accurately recapitulated (**Fig 2b, d**), including two hydrogen bonds to the side hydroxyl groups of CHD and a hydrogen bonding network around the CHD head hydroxyl group (CHD_r1, **Fig 2c**). This structure demonstrates that our interface design pipeline can accurately design detailed protein-ligand interactions.

**Figure 2.**
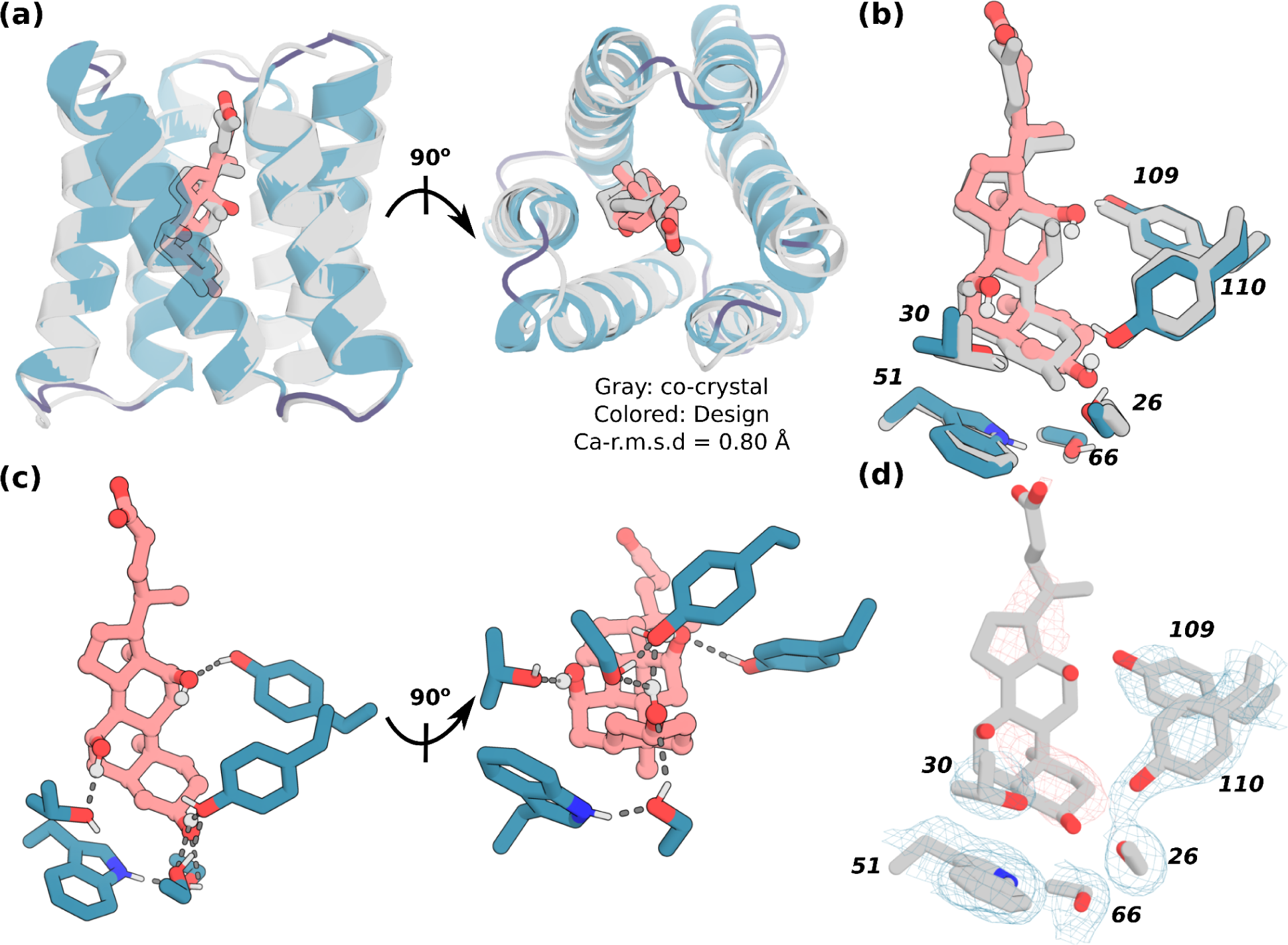
X-ray crystallography demonstrates accuracy of design approach. **(a)** The crystal structure of CHD_r1 (gray) is very similar to its computational design model (teal). **(b-c)** The designed sidechain interactions with the three CHD hydroxyl groups and the buttressing hydrogen bonding network are closely recapitulated in the crystal structure (design is colored, crystal structure is in gray). **(d)** The ligand and the key interacting residues were well resolved with clear electron density. The protein backbone is shown in cartoons, and CHD and the key interacting side chains in sticks. Pink, ligand carbon atoms; red, oxygen; blue, nitrogen; white, polar hydrogen. The residue numbers of the key residues are labeled. Also see **Fig S7a-c**.

### Generating higher affinity designs by scaffold resampling

We next sought to generate higher affinity designs by sampling new backbones around the pseudocycle scaffolds which gave rise to the first round hits. We generated ∼5000 sequences for each scaffold using proteinMPNN(*13*) and predicted their structures using AF2(*14*); this generates backbones which range from 0.5 to 3 Å Ca-r.m.s.d from the starting template (**Fig S1b**). The pre-generated ligand conformers were docked against the corresponding resampled scaffolds using RifDock(*3*) generating 10-30 million new docks. After two-step predictor selection, 50,000-500,000 docks were selected for PSSM-based Rosetta design protocol or ligandMPNN design protocols, as described above. 5,000-15,000 de novo designs **(Table S1)** were selected for a second round of binder screening (see “***High-throughput methods for binder identification”*** in Methods). As expected, the designs from the second round showed significantly higher SC to the ligand compared to the designs from the first round (**Fig S1c**).

We observed significant improvement in binder quality and quantity in the second round design round (**Table S1** & **S2**). For CHD, the highest affinity binders among all the 37 verified binders (as measured by FP) improved by ∼700 fold from 3.2 and 5.3 μM (**Fig 1c** & **S2a-b**) in the first round to 4.68 nM in the 2nd round (**Fig 3a-d** & see full binder list **S8-10**). Site saturation mutagenesis (SSM) of the five highest affinity 2nd round binders (CHD-d1 to d5) confirmed that the key interactions in the design model were essential for binding (**Fig S11**). For T44, the affinity by FP improved from 47.4 nM to 18.2 nM (**Fig 3g-h, S14**-**15**). For AMA and MTX, the binding affinities in the first round were too weak to be measured off the yeast surface (where there is very high avidity), likely in the high micromolar to millimolar range. In the second round, we obtained 870 nM binders (measured by surface plasmon resonance) for AMA (**Fig 3e** & **S12**, SPR, see ‘***SPR studies***’), and for MTX, the most challenging target (**Fig 1e & S6a**), we obtained a 6.9 μM binder (MTX-d1) from one of the β barrel-like scaffolds (**Fig 3f**), a first example of a de novo binder to a highly polar and flexible ligand (**Fig S4e** & **Table S4**). The designed binding mode of MTX-d1 is supported by SSM analysis and competition assays (**Fig S16**). As in the first round, binders were obtained using both the Rosetta-based and ligandMPNN-based interface design protocols (**Table S2**) but the ligandMPNN based method generated generally tighter binders. Control experiments for CHD and MTX in which the backbone and docking pose were fixed did not yield similar or any improvements in affinity(*16*), highlighting the importance of resampling for increasing shape complementarity and binding affinity.

**Figure 3.**
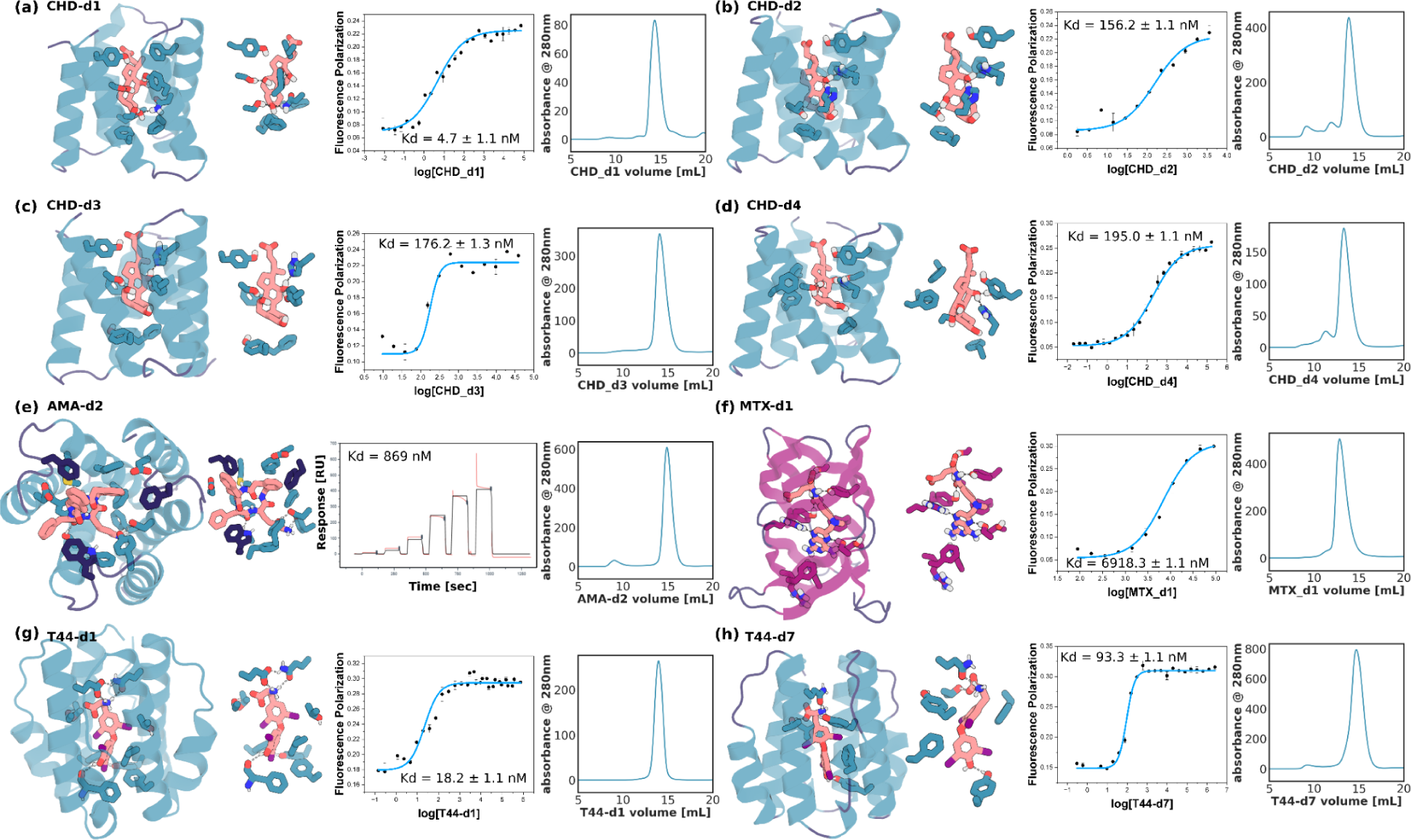
Experimental characterization of selected designed binders from the affinity improving round. **(a-d)** Nanomolar affinity CHD binders CHD_d1 to d4 (full list in **Fig S9**), **(e)** nanomolar binder for AMA (see **Fig S12**); **(f)** micromolar methotrexate binder, **(g-h)** two nanomolar T44 binders (full list at **Fig S14**). For each panel, from left to right, the design model, zoom in on the sidechain-ligand interactions, FP (or SPR in the case of AMA) binding measurements, and SEC traces. Kd values and error bars are from two independent experiments. Interacting side chains and ligands are shown in sticks, with oxygen, nitrogen, iodine, and polar hydrogen colored in red, blue, purple, and white, respectively. Key interactions are indicated by gray dashed lines. The cartoon of and sticks from helixes, sheets, and loops are colored in teal, magenta, and dark blue, respectively.

### Converting pseudocycle binders into sensors

Because in pseudocycles the stabilizing interactions are primarily within repeat units and between adjacent units, all of the designs by construction can be split into two or more chains that are likely to fold at least in part in isolation (**Fig 4a, g**, this contrasts with most globular proteins, where splitting is likely to disrupt the central hydrophobic core). Because of this, ligand binding can be coupled to reconstitution of the full pseudocycle.

**Figure 4.**
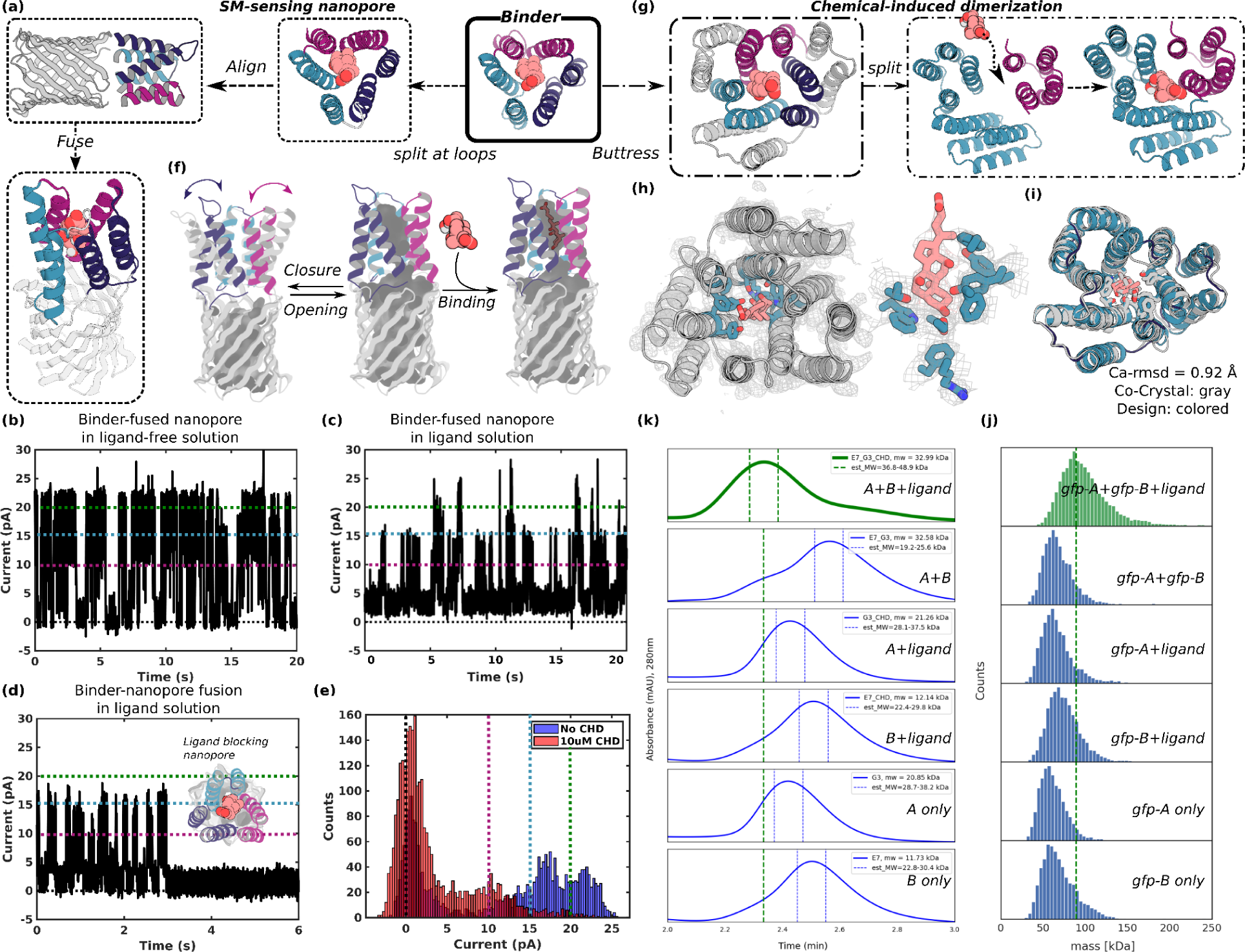
Conversion of pseudocycle binders into ligand gated channels and CID systems. **(a-f)** Ligand sensing de novo nanopore construction. **(a)** The three structural repeat units of CHD_r1 were inserted into three different loops in a 12 stranded de novo nanopore using inpainting to join the chains such that the central axes of binder and nanopore are aligned. The conductance of the original nanopore is ∼220 pS **(Fig S17a)**, and is not influenced by CHD. The conductance of binder-fused nanopore in the absence **(b)** and presence of CHD **(c-e)**: In the absence of CHD **(c)**, the pore fluctuates between a state with high conductance very similar to the unmodified pore and a low conductance state **(c)**; in the presence of CHD the duration of the low conductance states is greatly increased **(c-e)**; the longer record **(c)** and a single closure event **(d)** are shown for clarity, and the histogram of the current with and without ligand is shown in **(e)**. Different currents, 0, 10, 15, 20 pA are marked out for clarity using dashlines in black, magenta, teal, and green, respectively in **b-e**. The gated nanopores are robust through multiple cycles of opening and closure **(c)**, upon reversal of the voltage, the original high conductance state is restored **(Fig S17b)**. **(f)** The conductance fluctuations of the binder-fused pore in absence of ligand likely reflect transient association of the 3 subunits; ligand binding stabilizes the associated state leading to prolonged blocking of the pore. **(g-k)** CID system construction. CHD binder, CHD_r1, was buttressed by diffusion of an outer ring of helices to increase the stability of split protein fragments **(g** & **Fig S18**). The crystal structure of the buttressed binder with ligand **(h** & **Fig S7d-f**) is in close agreement with the design model **(i)**. To create a CID system, we split the buttressed binders into halves and redesign the protein-protein interface to increase solubility of the fragments and disfavor association in the absence of ligand. Characterization of CHD induced association of the split fragments by size exclusion chromatography **(k)** and mass photometry **(j)**. Dimerization of the two split domains (A and B) in presence (first trace from top), but not the absence, (second trace from top) of ligand. The individual monomers do not dimerize in the presence(third and forth trace from top) or absence (fifth and sixth traces from the top) of ligands. N terminal gfp tags were fused to the monomers to facilitate detection by mass photometry.

We first took advantage of this property by integrating a CHD binder, CHD_r1, consisting of 3 repeats of a helical hairpin around the central binding site, into a de novo designed 12 stranded beta barrel nanopore(*24*) such that reconstitution of the complete pseudocycle from the split domains would block ion conductance through the pore **(Fig 4a)**. We aligned the central axis of CHD_r1 with the central axis of the designed nanopore (TMB12_3)(*24*), and inserted the three helical hairpins of the binder between strands 3 and 4, 7 and 8, and 11 and 12 of the beta barrel using RosETTAfold joint inpainting(*25*) to build short connectors (5-8 Å) between the binder hairpins and the nanopore (**Fig 4a**, see **‘*Design of SM binder fused nanopore’*** in Methods).

We expressed 12 integrated CHD binder-nanopore fusions in *E. coli* and purified them from inclusion bodies (see “***Design of SM binder fused nanopore”*** in Methods). Five designs with monomeric SEC peaks were integrated into planar lipid bilayers, and ionic conductances were measured under an applied voltage of 100 mV. The open-pore conductance of the original nanopore is constant at ∼220 pS and was not affected by CHD (**Fig S17a**). Two of the binder-nanopore fusions had similar open-pore conductances (**Fig 4b**) to the original nanopore (**Fig 17Sa**), but showed frequent transitions to two lower conductance states which likely reflect transient association of 2 or 3 helical hairpins in the absence of the ligand (**Fig 4f** left). In the presence of CHD, the dwell time of the nanopore in the closed state significantly increased for the most sensitive design (**Fig 4c-e**), likely reflecting stabilization of the trimeric pseudocycle by the ligand (**Fig 4f** right). The current amplitudes approached near zero for ∼0.4-3 seconds on average before returning to higher conductance (**Fig 4c-e**); each fluctuation to a low conductance state likely corresponds to an independent CHD binding event indicating that the pores are robust to multiple cycles of CHD binding and release (**Fig 4c**). The limited number of observed states enabled quantification of the pore blocking and release dwell time distributions, which were consistent with the binding affinity of the unfused CHD binder (**Fig S17c-d**). In some cases the blocked state was very long lived (**Fig 4d**), switching back to fluctuating between open and closed states only upon reversal of the applied voltage polarity without compromising the stability or the conductance of the nanopore (**Fig S17b**). The lower noise and considerable increase in duration of the closed state conductances at longer timescales (100 ms to s) in the presence of the ligand (the fraction of transitions to closed states with dwell times of one second or more increased from 1% to 6.5% upon addition of CHD) leads to lower sampling frequency requirements compared to previously engineered sensors(*26*–*28*). The simplicity of construction of nanopore sensors by incorporating designed pseudocycles binding targets of interest into de novo designed quiet nanopores should enable the generation of a wide variety of new sensors with superior properties.

We next sought to convert the pseudocycle binders into chemically-induced dimerization (CID) systems (**Fig 4g**). There has been considerable interest in CID systems in protein engineering and synthetic biology for SM inducible switches for regulating protein association(*29*), signaling or enzymatic activity(*30*). Despite the potential, almost all work has utilized a small number of CID systems, such as rapamycin-FKBP-FRB(*31*), and involved natural proteins as one(*32*) or both partners(*33*). Synthetic biology approaches would benefit considerably from systematic approaches for designing CID systems for new ligands, for example for feedback control based on product levels in metabolic engineering.

We first stabilized the pseudocycles by building a second ring of stabilizing structural elements around the inner ring which forms the binding interface (**Fig 4g** & **S18a**): the interactions between the inner and outer rings should contribute sufficient stabilization to make up for the loss in interactions between the N and C terminal repeats upon splitting. We chose the CHD binder, CHD_r1, as a proof-of-concept, and incorporated a second ring of helices using RFDiffusion(*10*); the inner and outer helices packed closely around a hydrophobic core (**Fig S18a**). We obtained a co-crystal structure of the buttressed binder with CHD bound (**Fig 4h,i** & **S7d-f**) with the outer buttressing ring of helices, the ligand conformation and key interacting sidechains very similar to the design model (**Fig 4h-i** & **S7e-f**). The buttressed binder was more stable and bound to CHD more tightly, likely due to rigidification of the target binding conformation (**Fig S18b**). This approach of testing binding first at the single layer stage and then adding an outer layer by buttressing to stabilize the highest affinity binders has the considerable advantage of reducing the difficulty and cost of design testing: the single ring structures are less than 120 aa and can be encoded on two 250-300 nt commercially available oligonucleotides from an oligo array, whereas the two ring structures are longer than 200aa for which gene synthesis is far more expensive and lower throughput.

To generate a CID system, we split the buttressed binder into two parts (**Fig 4g**). Using tied-position proteinMPNN design (see “***CID design and testing”*** Methods), we optimized the newly created interfaces to increase the solubility of the split domains while retaining the ability to form the holo-complex in the presence of the ligand. We expressed, purified, and tested 34 designed CID protein pairs and determined their oligomerization state by SEC (**Fig 4k**) and mass photometry (**Fig 4j**) in the presence and absence of CHD. For the best behaving of these designs, the individual domains were monomeric in isolation but associated to form the heterodimer in the presence of 10 uM CHD in both experiments. Because the individual subdomains (A is 9.8 kDa, B is 18.9 kDa) of our designed CID system are smaller than the detection limit of mass photometry (around 30 kDa), we expressed A and B with a *N*-terminus green fluorescent protein (gfp). By both SEC and mass photometry, dimerization was observed with ligand (**Fig 4j**, first lane from top) but not without ligand (**Fig 4j**, second lane from top). Individual proteins with (**Fig 4j**, third and forth lane from top) or without (**Fig 4j**, fifth and sixth lane from top) ligands did not show dimerization. The mass of the dimer peak derived from mass photometry, 89.0 ± 22.0 Da, was close to the expected 83 kDa for the gfp-A-ligand-gfp-B complex. As protein association can be readily coupled to site specific transcriptional activation (for example by linking a DNA binding module with an activator module) and other cellular readouts, the ability to generate CID systems for potentially any SM of interest could have very broad impact for metabolic pathway engineering by enabling feedback control based on levels of the desired product and/or unwanted or intermediate species.

## Conclusion

Our SC pseudocycle-based design approach goes considerably beyond previous design studies(*3*–*9*) in generating binders to large, polar, and/or flexible SMs, such as MTX, T44, and AMA, with affinities in the range for use in diagnostics without experimental affinity maturation. The crystal structures of the CHD binders with the intricate network of side chain-ligand interactions nearly identical to the computational design model highlights the accuracy of the design approach. While the importance of SC for binding has been long understood(*1*), our direct optimization of SC by design would have been very difficult prior to the recent advances in DL-based protein structure prediction and design(*13*–*15*), which enabled rapid de novo sampling and evaluation of pseudocycle topologies(*11*), resampling of the best solutions for a given ligand, and design of interactions with the ligand (with the new LIgandMPNN); the ability to iteratively resample the backbones and docking poses was essential for binding affinity improvement.

Our pseudocycle-based approach has a number of advantages for ligand binder and sensor design. First, centrally located and high SC binding pockets can be generated with high affinity for very diverse SMs. The variety of possible individual repeat unit structures is almost unlimited, and the number of repeat units can be readily varied, leading to vast numbers of possible pseudocycle structures all harboring central pockets. Because the structures are largely locally encoded with few long range contacts, interactions with each portion of the ligand can subsequently be optimized independently. Second, the ability to test small genetic footprint single rings forming the binding pocket in a first step, and then buttress the best designs in a second step, enables rapid and low cost gene synthesis for exploring diverse design solutions without compromising the robustness and stability of the final binding modules. Third, because of the local encoding of the structure, designed binders can be readily integrated into ligand gated channels and CID systems by splitting into subdomains that retain most of the interactions present in the full protein. The power of de novo design is highlighted by our ability to integrate the three repeat units of the CHD binder into three loops of a robust designed nanopore to create a CHD gated nanopore with almost complete gating of current by CHD; because of the high signal to noise ratio, minimal post processing of the acquired signal is required compared to previously engineered pores based on native proteins, which typically exhibit high level of noise in the absence of ligand due to the presence of a multitude of partially occluded states(*26*, *34*). Here we used RFdiffusion(*10*) to build the outer pseudocycle to buttress the inner ring; moving forward it should be possible to adapt RFdiffusion All-Atom(*9*) to generate single ring pseudocycles directly around target ligands. We anticipate that our SC pseudocycle-based approach should enable generation of robust ligand responsive channels and sensors for a wide variety of molecules of biological interest.

## Acknowledgments

We thank Dr. Sam Pellock and Dr. Indrek Kalvet for providing many general discussions for laboratory settings. We thank Avi Swartz and Dr. Patrick Erickson for providing chromatography assistance. We thank Dr. Buwei Huang and Dr. Robert Ragotte for providing SPR assistance. Crystallographic diffraction data was collected at the Northeastern Collaborative Access Team beamlines at the Advanced Photon Source and at CBMS/NSLS2. NECAT is funded by the National Institute of General Medical Sciences from the National Institutes of Health (P30 GM124165). This research used resources of the Advanced Photon Source, a U.S. Department of Energy (DOE) Office of Science User Facility operated for the DOE Office of Science by Argonne National Laboratory under Contract No. DE-AC02-06CH11357. The Center for Bio-Molecular Structure (CBMS) is primarily supported by the NIH-NIGMS through a Center Core P30 Grant (P30GM133893), and by the DOE Office of Biological and Environmental Research (KP1607011). NSLS2 is a U.S.DOE Office of Science User Facility operated under Contract No. DE-SC0012704. This publication resulted from the data collected using the beamtime obtained through NECAT BAG proposal # 311950. Dr. Meerut Said, Long Tran, and Dr. Sgardip Majumder are co-second authors, they contributed equally, and everyone agrees their authorship orders can be exchanged to benefit their own career development.

## Authors contributions

L.A. designed the project, designed and wrote the scripts of the whole computational pipeline, designed and performed the first round of screening for ligand CHD, MTX, T44, designed and performed the second round of screening for ligands CHD, MTX, AMA, T44. L. T., L.A., and D.J performed the buttrase of the binders. L.T. and L.A. performed the design and test of the CID. L.T. helped manuscript preparation. M.S. designed and performed the first round of screening for ligand AMA, assembled some of the yeast surface display libraries using qPCR. S.M. instructed V.A. with L. A., and designed, tested the binder-nanopore fusion with assistance from V.A. S.M. also wrote the nanopore part of the manuscript. I.G. led and assembled some of the yeast surface display libraries using qPCR, transformation of all yeast surface display libraries, and all NGS experiments. G.R.L. wrote the prototype of the repetitive ligandMPNN design script, and the ligand tail detection script. J.D. provided access to the ligandMPNN script and many general discussions. I.A. provided access to the ChemNet script and many general discussions. B.C. provided the prototype and ideas for two-step predictor scripts, many help on scripts and many general discussions. P.S. assisted S.M. on designing of the cyclic peptide ligand. A.B. generated the model of the co-crystal structure. A.K. performed all the crystal screening work. A.P. and L.A. performed the mass photometry assay. D.Z, D.R.H. and L.A. wrote the protein pocket detection scripts, optimized the proteins for docking through examining the proteins, and permuted the proteins to adjust the N, C terminus. D.F. wrote the NGS assembly and matching script. C.N. proposed the idea of PSSM-based binder design, provided the prototype script for generating PSSM, and provided many script help during the early stage. X.L. performed all the mass spec verification of the proteins. P.M.L. performed verification and synthesis of the labeling of some of the ligands. M.G. and D.K.V. provided some yeast library sorting and NGS experiments under the leadership of I.G. A.A.V. and S.M. designed the de novo nanopore used for binder-nanopore fusion. D.B. and L.A. designed the project together.

## Data and Materials availability

All Data and scripts are available online. All scripts for stepwise sampling are available at github (repository will be public after BiorXiv release): https://github.com/LAnAlchemist/Pseudocycle_small_molecule_binder. All scripts for CID design are available at github https://github.com/iamlongtran/pseudocycle_paper (repository will be public after BiorXiv release). All sequencing data were analyzed using an in-house script, and the git repository is at: https://github.com/feldman4/ngs_app.

---

**Figure S1.**
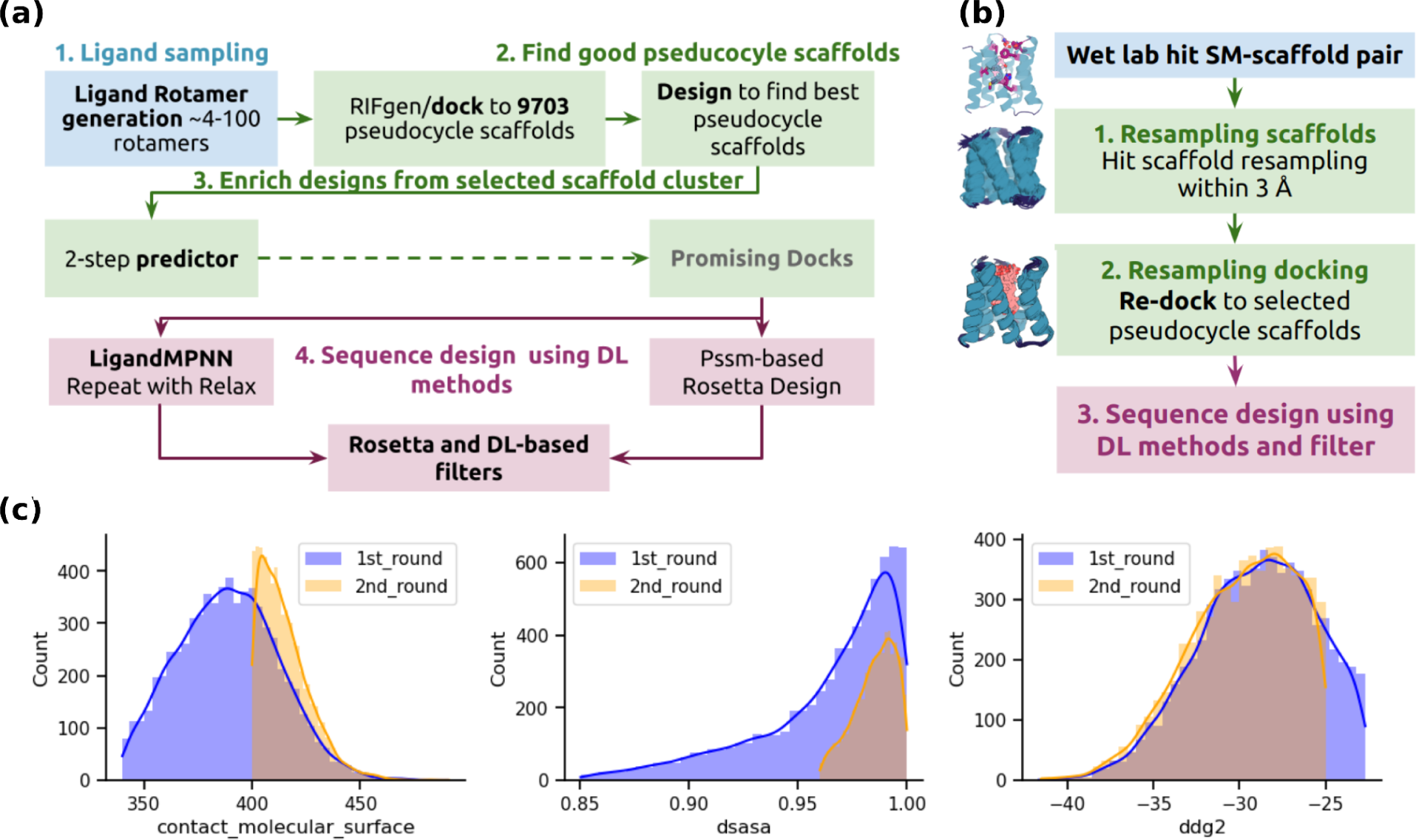
Computational design pipeline. **(a)** The flowchart of the first computational step to identify scaffolds suitable with the each SM ligands; **(b)** The flowchart of the second computational step to focus on the verified suitable scaffold-ligand pair and sampling the docking and the interface packing of the designed binder proteins; **(c)** The histogram plots comparison of MTX binders for order from the first step sampling (1st_round) v.s second step sampling (2nd_round). The contacts between the ligand and the protein got significant improvement through stepwise sampling.

**Figure S2.**
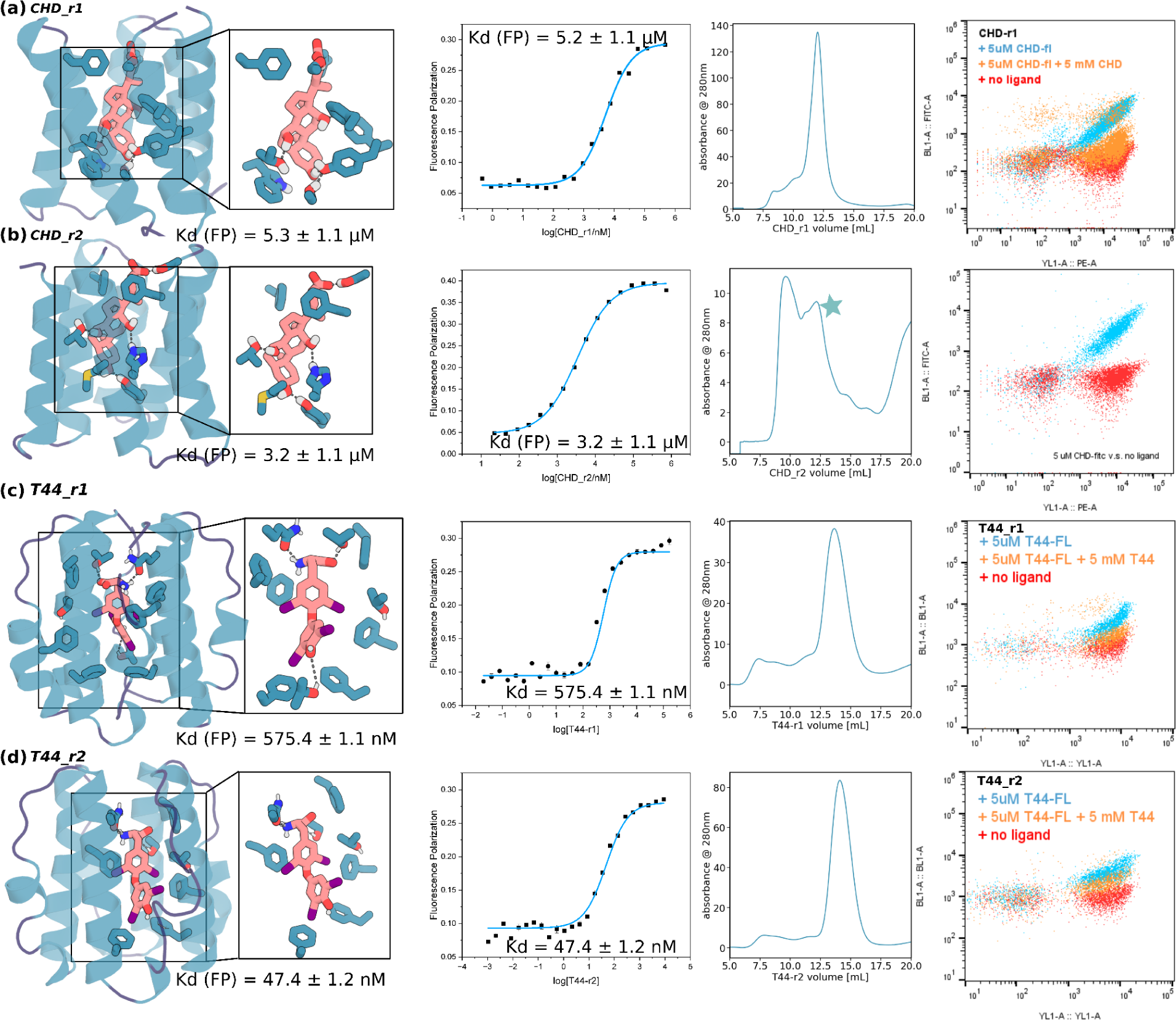
Characterization of the CHD and T44 binders from the first round of sampling. The initial hits from the first step screening are shown in each panel, the design model, interface, binding and competition assays from FP studies of the Two binders for CHD **(a-b)** and T44 **(c-d)**. The monomeric fraction of CHD_r2 was collected and marked with a star on the third panel in **(b)**. The designs were shown in cartoons, the ligand and key interacting residues were shown in sticks. Oxygen, nitrogen, and sulfur were colored in red, blue, and yellow, respectively.

**Figure S3.**
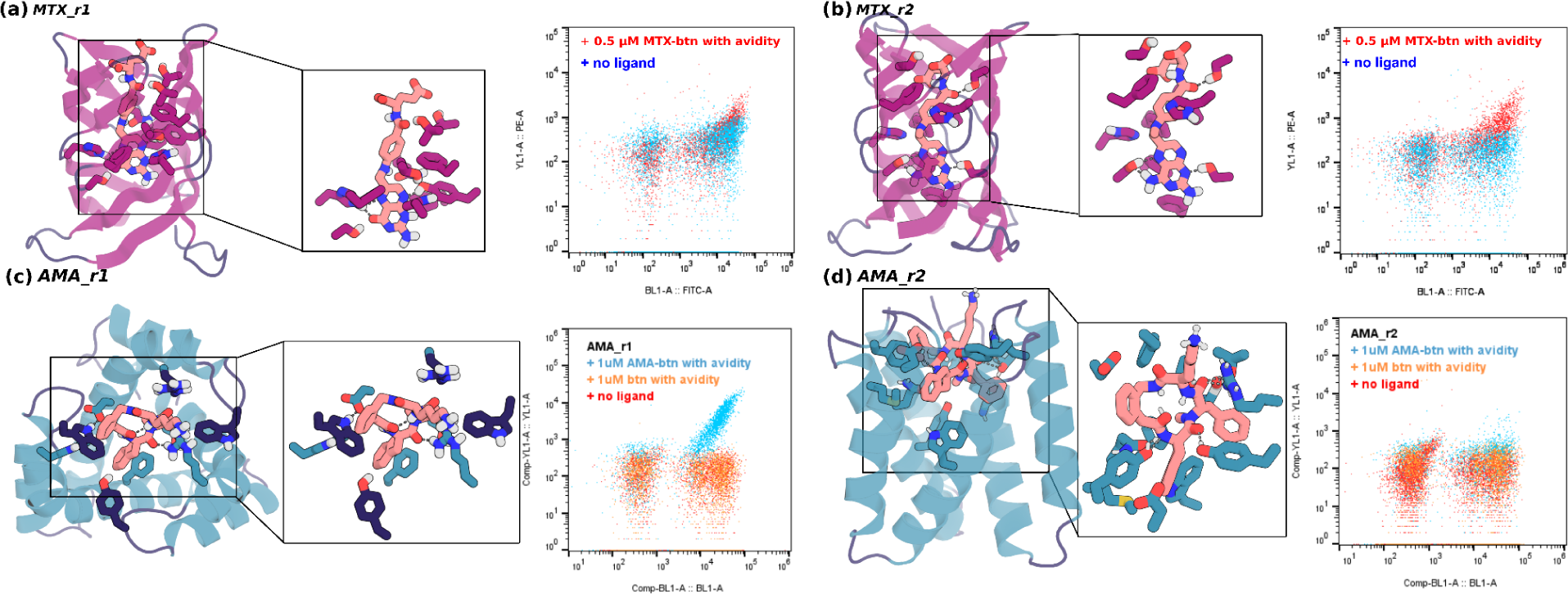
MTX and AMA binders from the first round of sampling. The initial hits from the first step screening are shown in each panel, the design model, interface, binding and competition assays from yeast surface display studies of the MTX **(a-b)** and AMA **(c-d)**. The binder expressed yeast showed a positive FITC signal (BL1-A), and the binding yeast showed a positive PE signal (BL1-A). The binder-expressing yeast showed clear strong double positive signals (red for MTX, blue for AMA), and only FITC signals when no biotin-labeled ligand presence (blue for MTX, orange for AMA). The designs were shown in cartoons, the ligand and key interacting residues were shown in sticks. Oxygen, nitrogen, and sulfur were colored in red, blue, and yellow, respectively.

**Figure S4.**
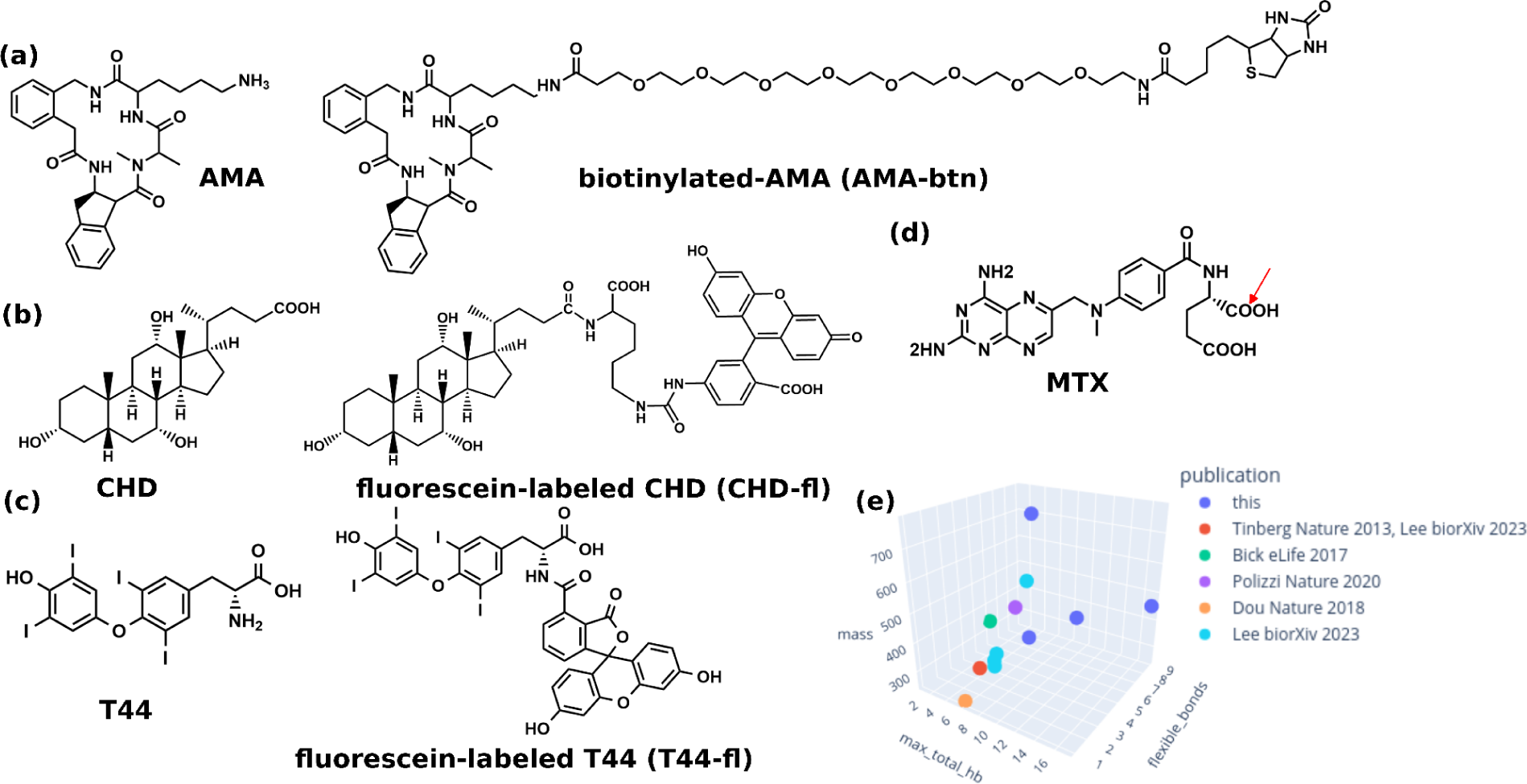
Target ligands. **(a-d)** The structure and the tagged version of AMA (synthesized by WuXi Chemistry Service), CHD (both commercially available), T44 (tagged version synthesized in house, no-tagged version commercially available), and MTX (commercially available for tagged and untagged). While both the tagged structure of fluorescein-labeled MTX (MTX-fl), and biotinylated MTX (btn-MTX) is commercially available, the structure was not reported, but the tagged site was reported, marked in red. **(e)** The features of ligands designed against in this work and previous works. Only ligands with affinity (Kd) and validation (SSM, binding competition, or crystallography) data are included. More details of ligand features, see **Table S4**.

**Figure S5.**
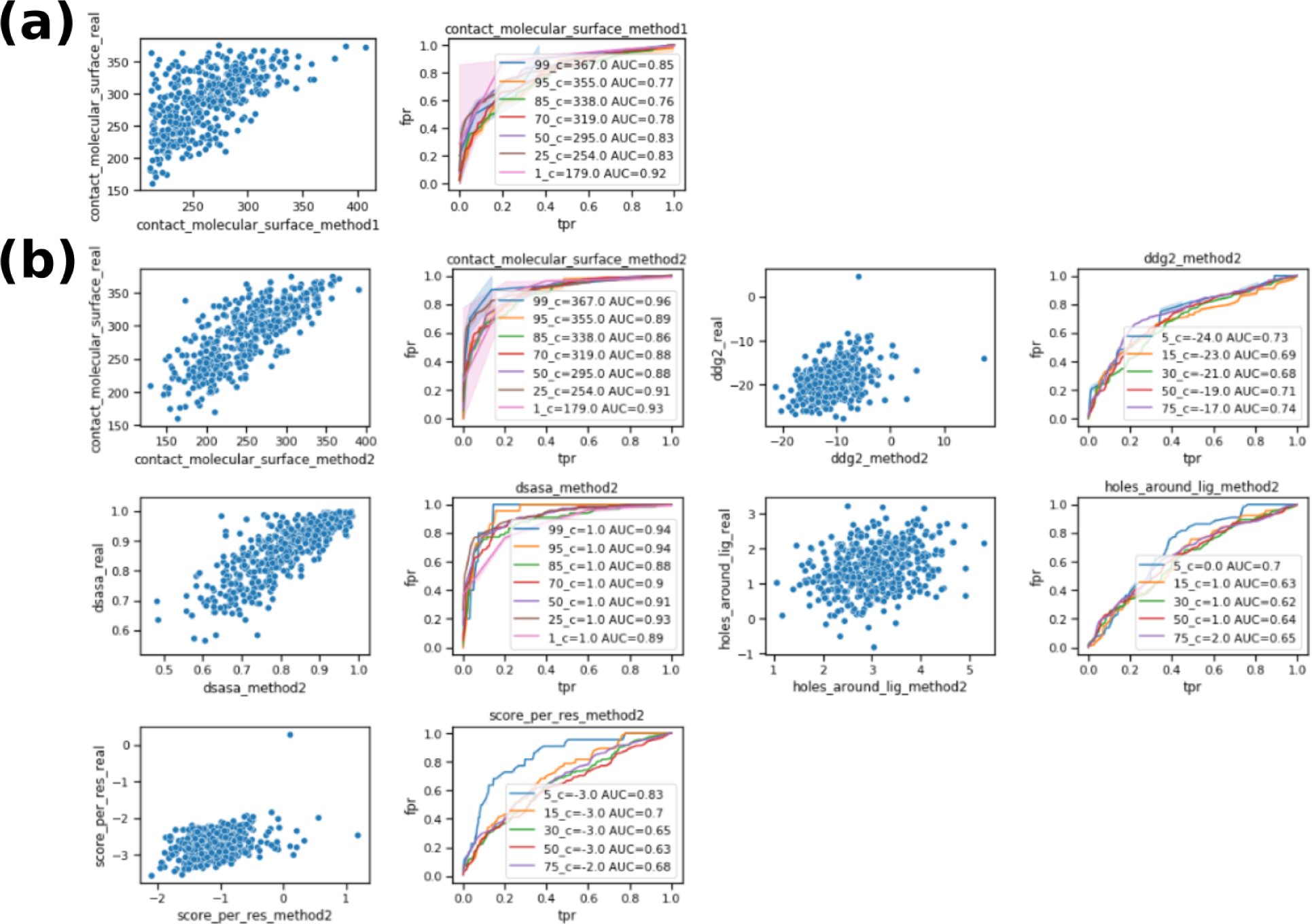
Two-step predictor scores are correlated with scores after full design. One example of evaluation on how predictive the two-step predictor is. 1000 docks to CHD were randomly selected to evaluable how the predictor works. For each set, the left panel is the scores generated from score from predictor (x-axis) v.s. final PSSM-based Rosetta design protocol (y-axis). The right panel is the receiver operating characteristic curve (ROC) curve at different quantiles to measure the predictiveness of the predictor script on the reported metric. The higher the false positive rate (fpr) with the increase of true positive rate (tpr), the more predictive the scripts are on the measured metric. The predictiveness at each quantile was also reported using area under curve (AUC), and the higher the AUC, the better the predictions are. **(a)** The CMS of input docks were clearly predicted well with predictor1; **(b)** The ‘CMS’, binding energy (‘ddg2’), burial of the ligands (‘dsasa’ and ‘holes_around_lig’) and the protein quality (‘score_per_res’) were highly predicted using the second predictor.

**Figure S6.**
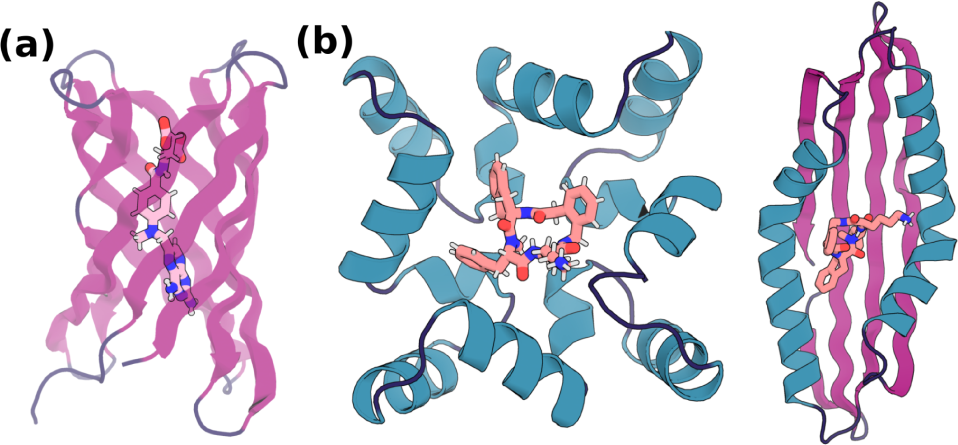
The other potential binding scaffold hits (not shown in. Fig 1**) from the first step sampling.** During first step HTP binder screening, for MTX and AMA, we also identified other scaffolds (apart from those shown in Fig 1b), which NGS enrichment suggested to be potential binders. These scaffolds were also identified as scaffolds in hit ligand-scaffold pairs and were used for second step computational sampling. Individual binders from these pseudocycle scaffolds were not verified individually for prioritization of other better performing binders. **(a)** One other potential binding pseudocycle scaffolds for MTX; **(b)** Two other potential binding pseudocycle scaffolds for AMA.

**Figure S7.**
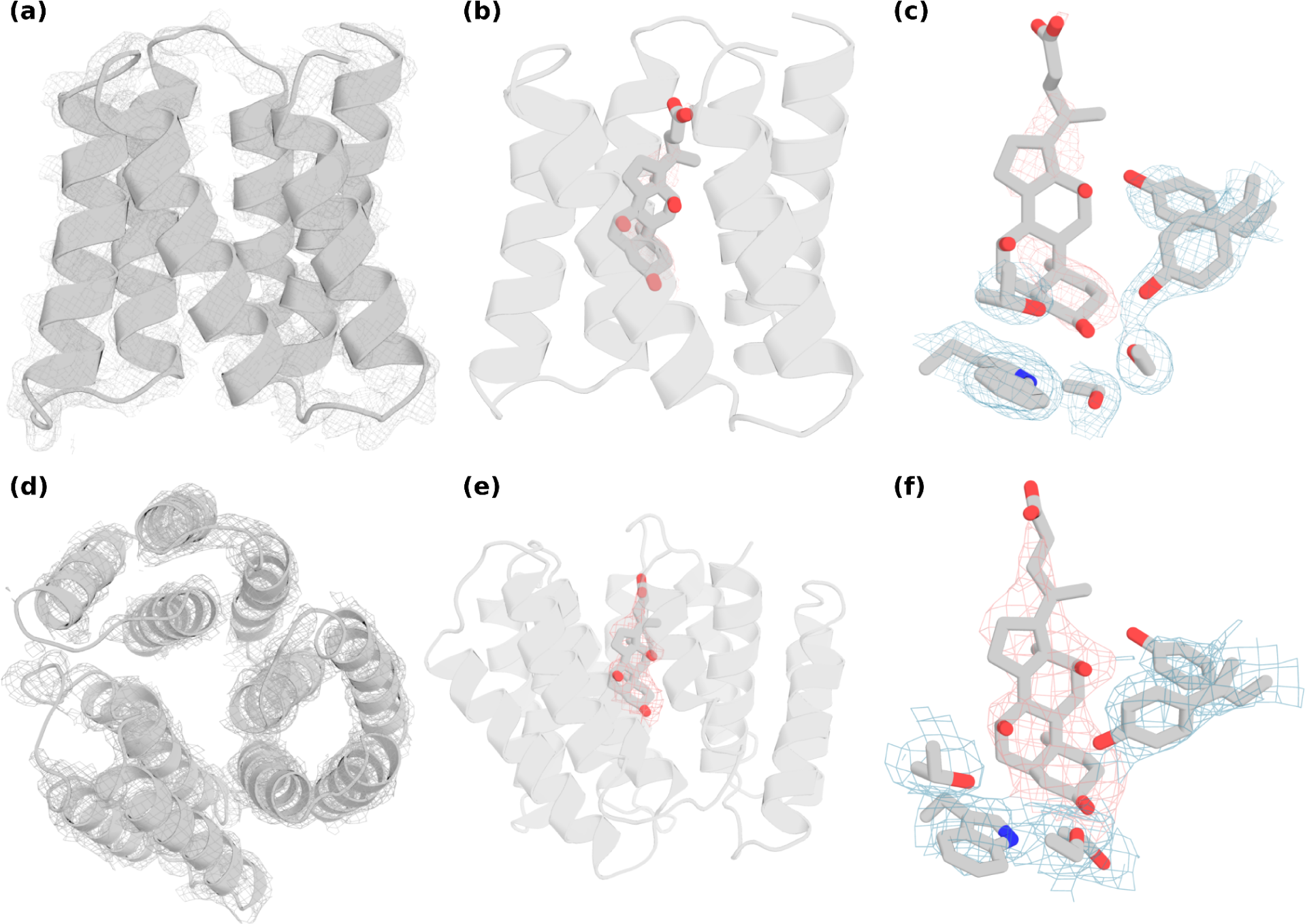
The comparison of crystal structures of CHD_d1 complex and CHD_buttress complex. The crystal structure of CHD_d1 **(a-c)** and CHD_buttress **(d-f)**. Only the protein backbone densities are shown in **a** and **d** for clarity. The backbone electron density, the ligand density, and the key interacting rotamer electron densities are shown from left to right. The electron density is shown in mesh and the refined co-crystal structure is shown in gray. The ligand and the key interaction rotamers are shown in sticks, where oxygen, nitrogen are colored in red and blue, respectively.

**Figure S8.**
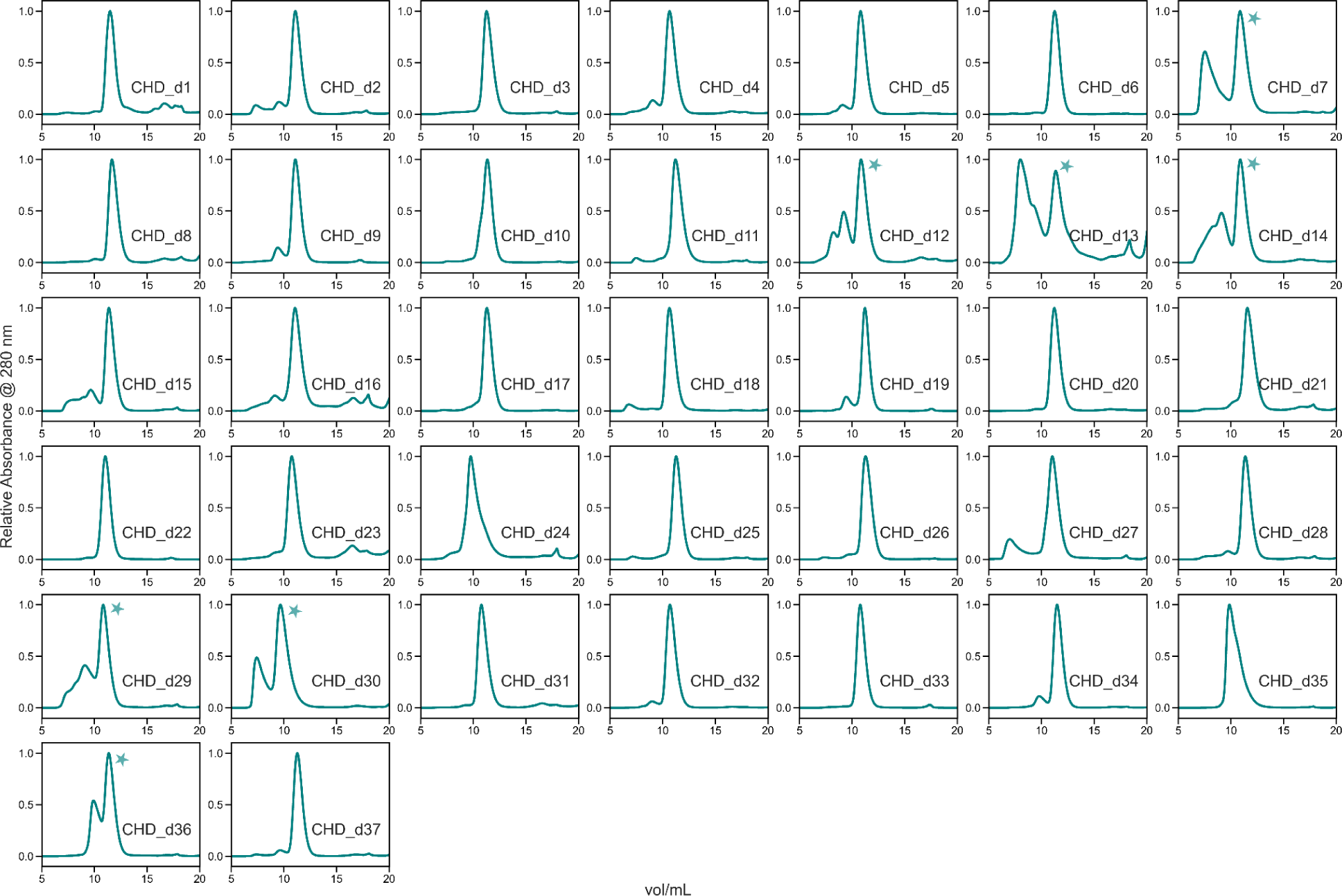
SEC profiles of 2nd round CHD binders. SEC traces of all CHD binders chosen for off-yeast binding test are shown here. For traces where multiple oligomeric states are present, the monomeric peaks collected for FP studies are marked with stars.

**Figure S9.**
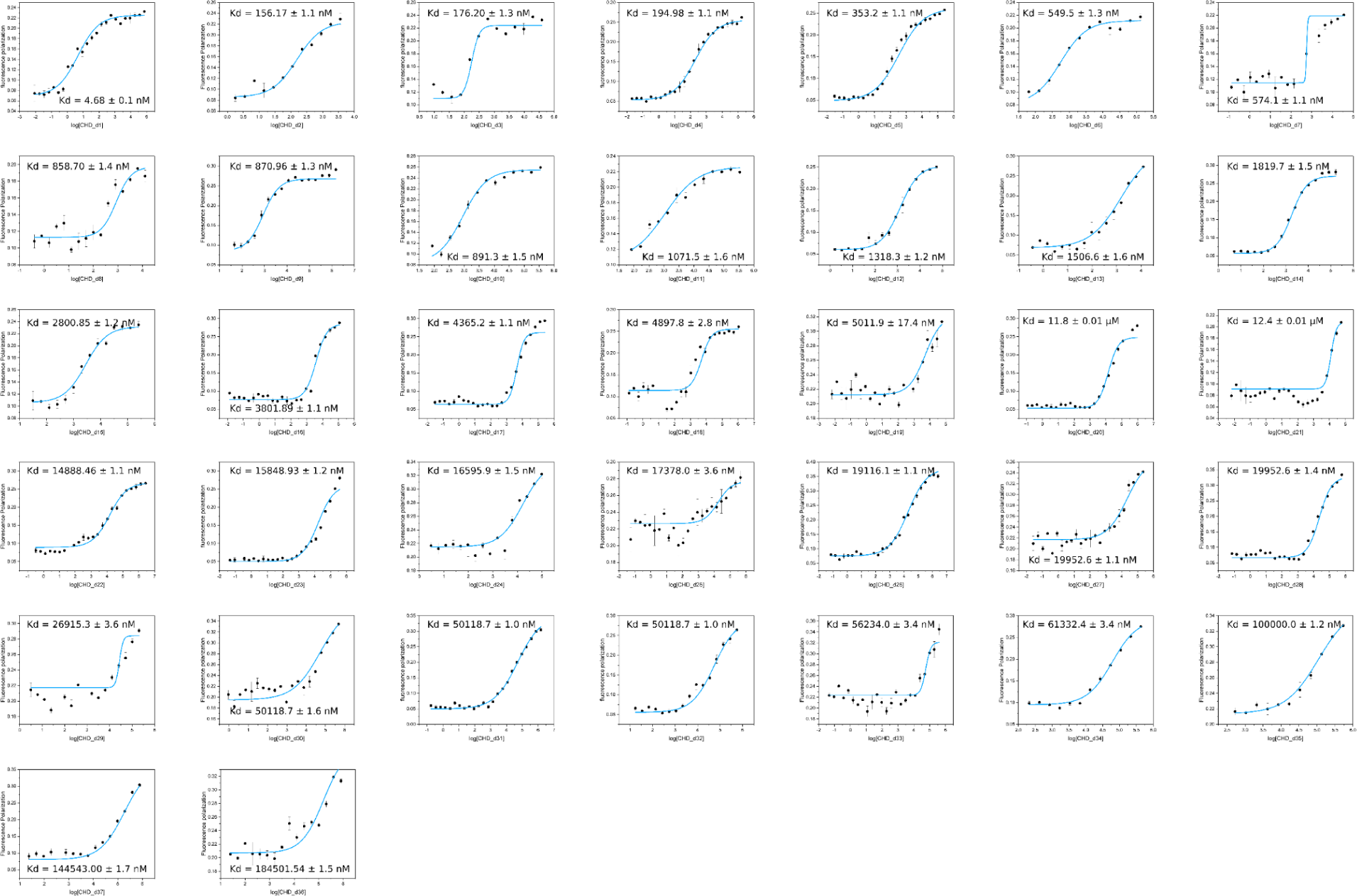
Fluorescence polarization titrations for all 2nd round CHD binders. FP profiles for all 37 purified 2nd round CHD binders; also see Fig 3.

**Figure S10.**
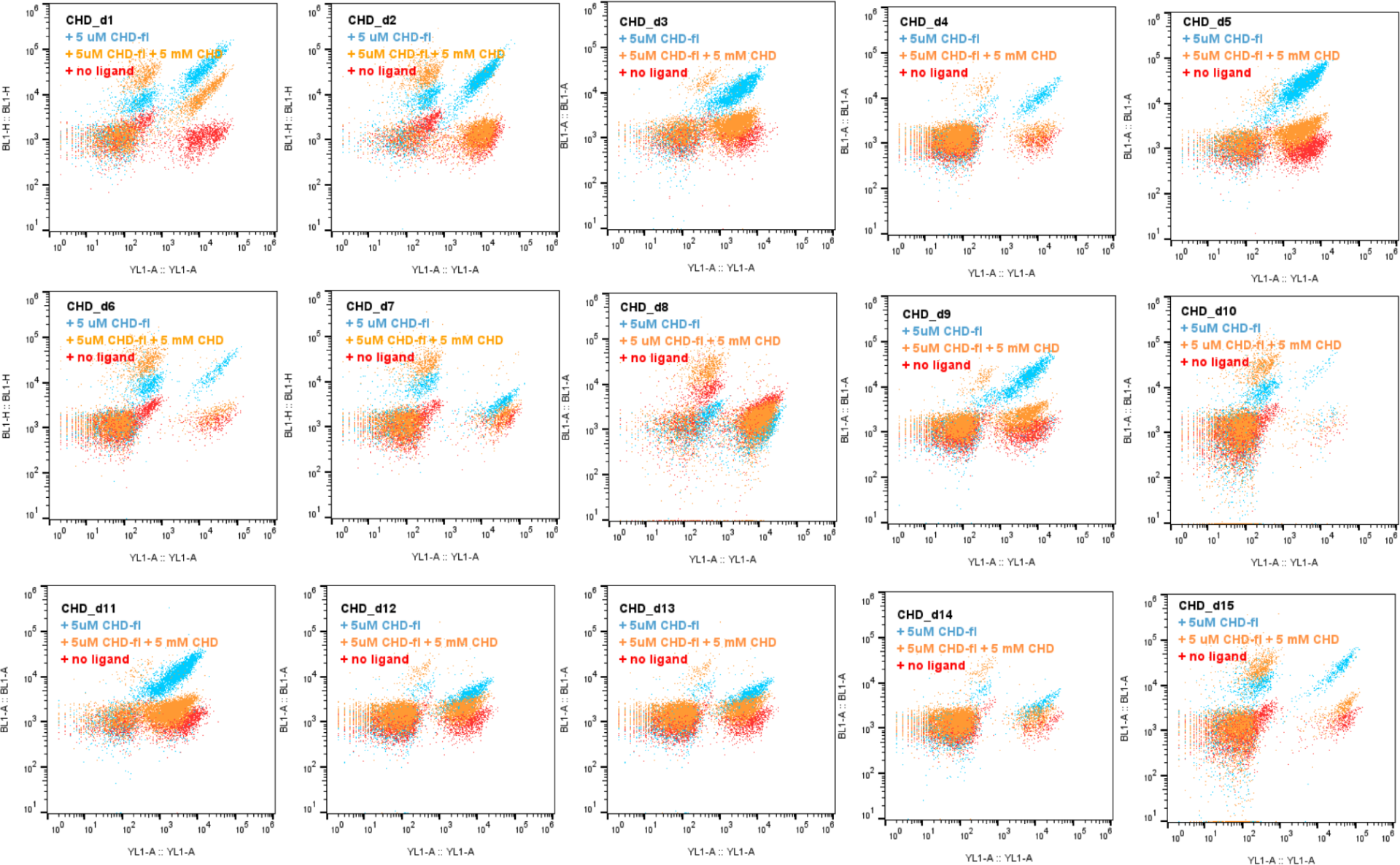
Competition of 2nd round CHD binders on yeast surface by free CHD. The designs were displayed on the surface of yeast, using 5 μM CHD-fl for binding and 5 mM free CHD for competition. The binder expressed yeast showed a positive PE signal (YL1-A), and the binding yeast showed a positive FITC signal (BL1-A). All binder expressed yeast showed clear strong double positive signals (blue), while weaker FITC signal upon free ligand competition (orange), and only PE signals when no FITC-labeThe binder expressed yeast showed a positive PE signal (YL1-A), and the binding yeast showed a positive FITC signal (BL1-A). All binder expressed yeast showed clear strong double positive signal (blue), while weaker FITC signal upon free ligand competition (orange), and only PE signal when no FITC-labeled ligand presence (red).

**Figure S11.**
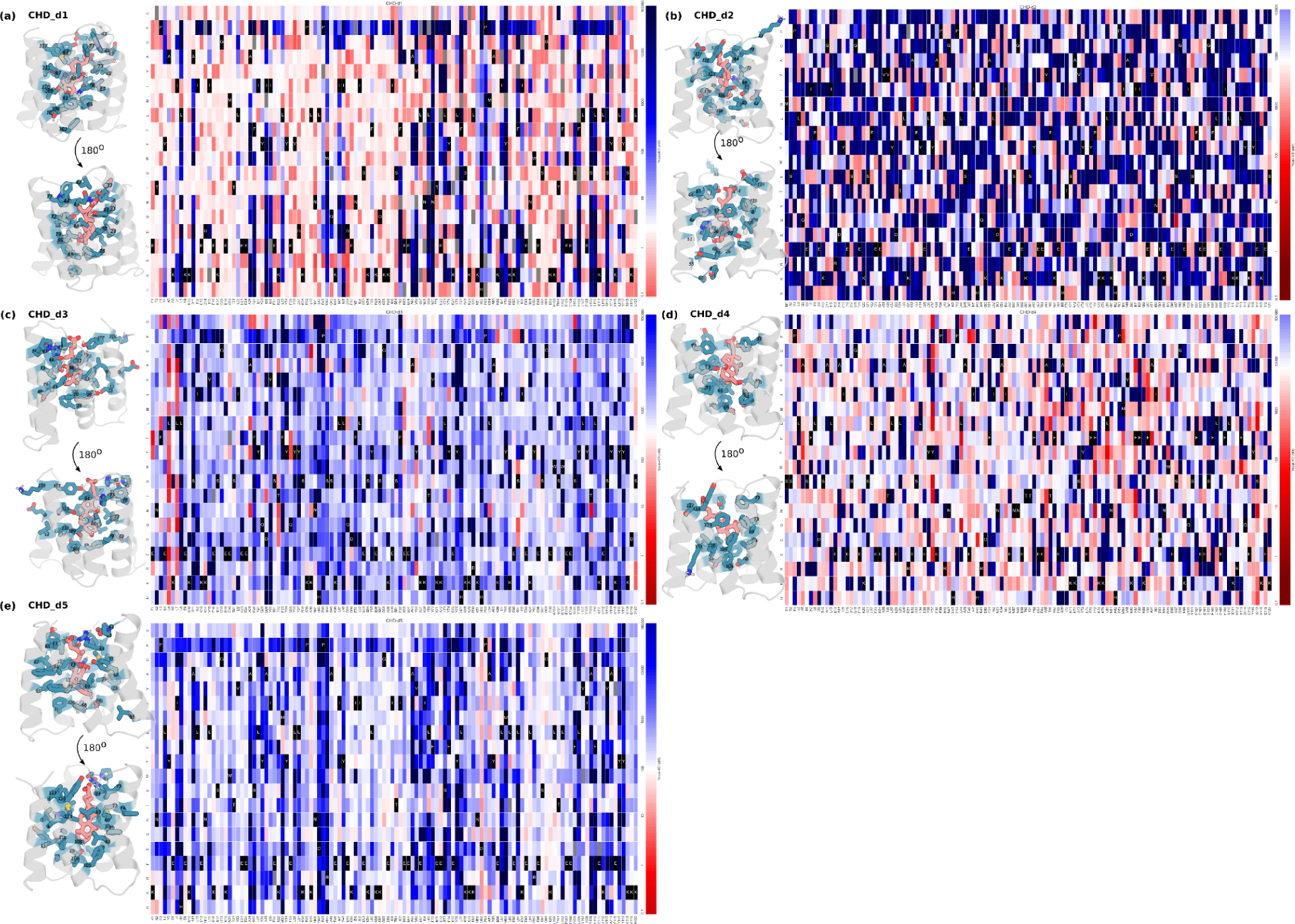
site saturation mutagenesis of CHD-d1 to CHD-d5. SSM analyses were performed for CHD-d1 to CHD-d5 (**a-e**). The interface residues were analyzed, and those conserved were colored in teal. All SSMs showed high conservation are the residues having contact with the ligand, supporting the designed conformations. The native residues are listed on the x axial, and colored in black. The estimated binding affinity of each mutant is colored based on y axial; the residue choices which may improve binding are more red, while the residues jeopardizing the binding are more blue. The conserved interface residues are colored in teal.

**Figure S12.**
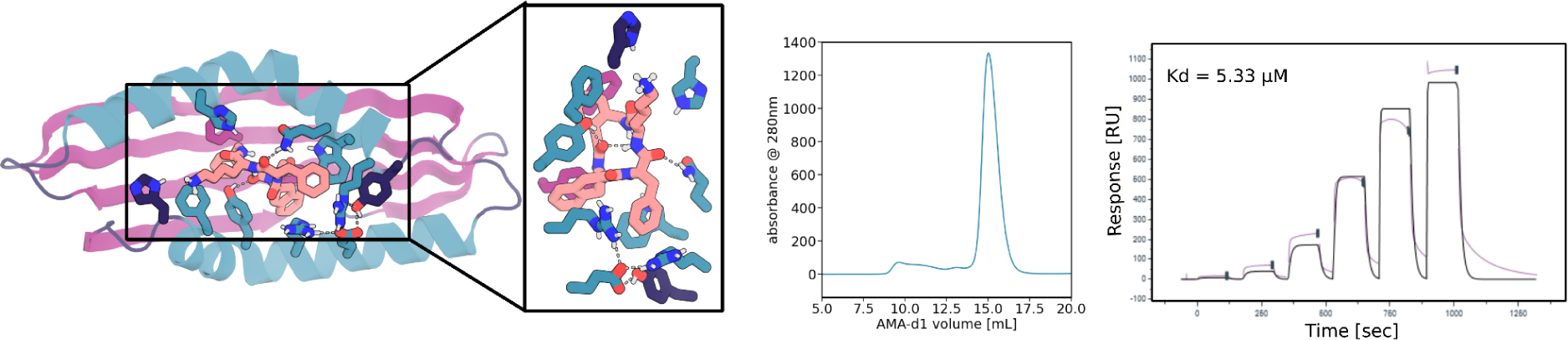
SPR analysis of additional 2nd round AMA binders not shown in. Fig 3. From left to right panel, the design model, SEC traces, and the SPR binding traces of AMA-d1 are shown in each panel. The design models are shown in cartoons, the ligand and the key interacting residues are shown in sticks. Oxygen, nitrogen are colored in red and blue, respectively. The cartoon of and the residues from helix, sheet, and loops are colored in teal, magenta, and dark blue, respectively.

**Figure S13.**
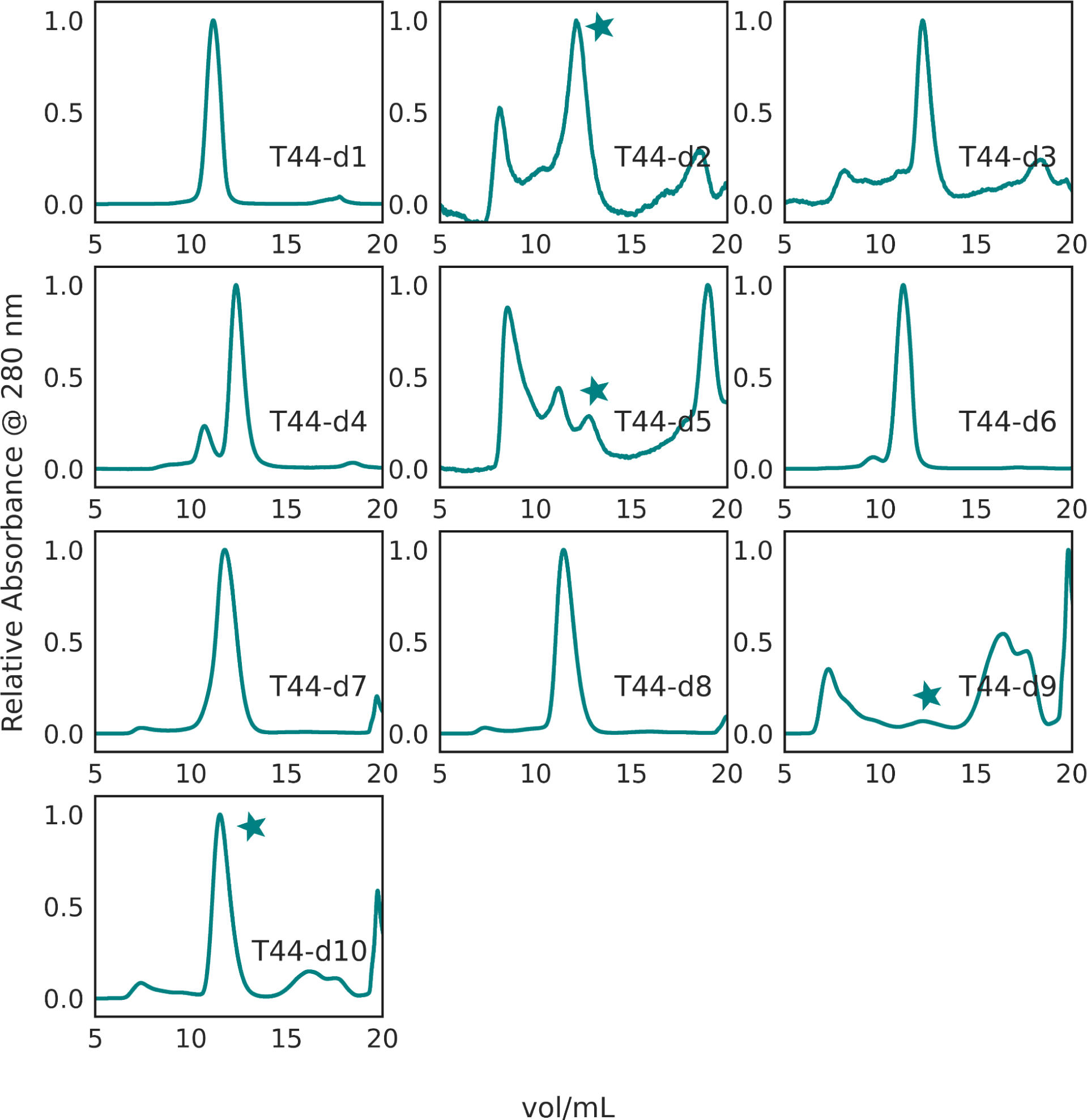
SEC analysis of second round T44 binders. SEC traces of T44 binders chosen for off-yeast binding test from the second round of sampling are shown here. For traces where multiple oligomeric states are present, the monomeric peaks collected for FP studies are marked with stars.

**Figure S14.**
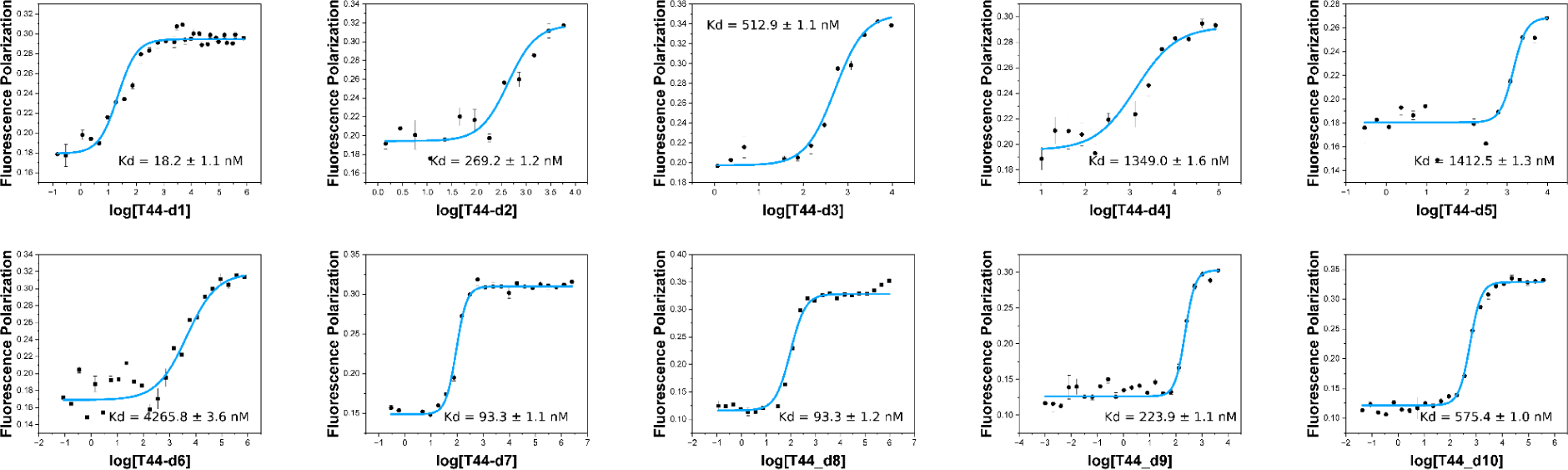
FP analysis of 2nd round T44 binders. FP profiles for all 10 purified 2nd round T44 binders; also see Fig 3.

**Figure S15.**
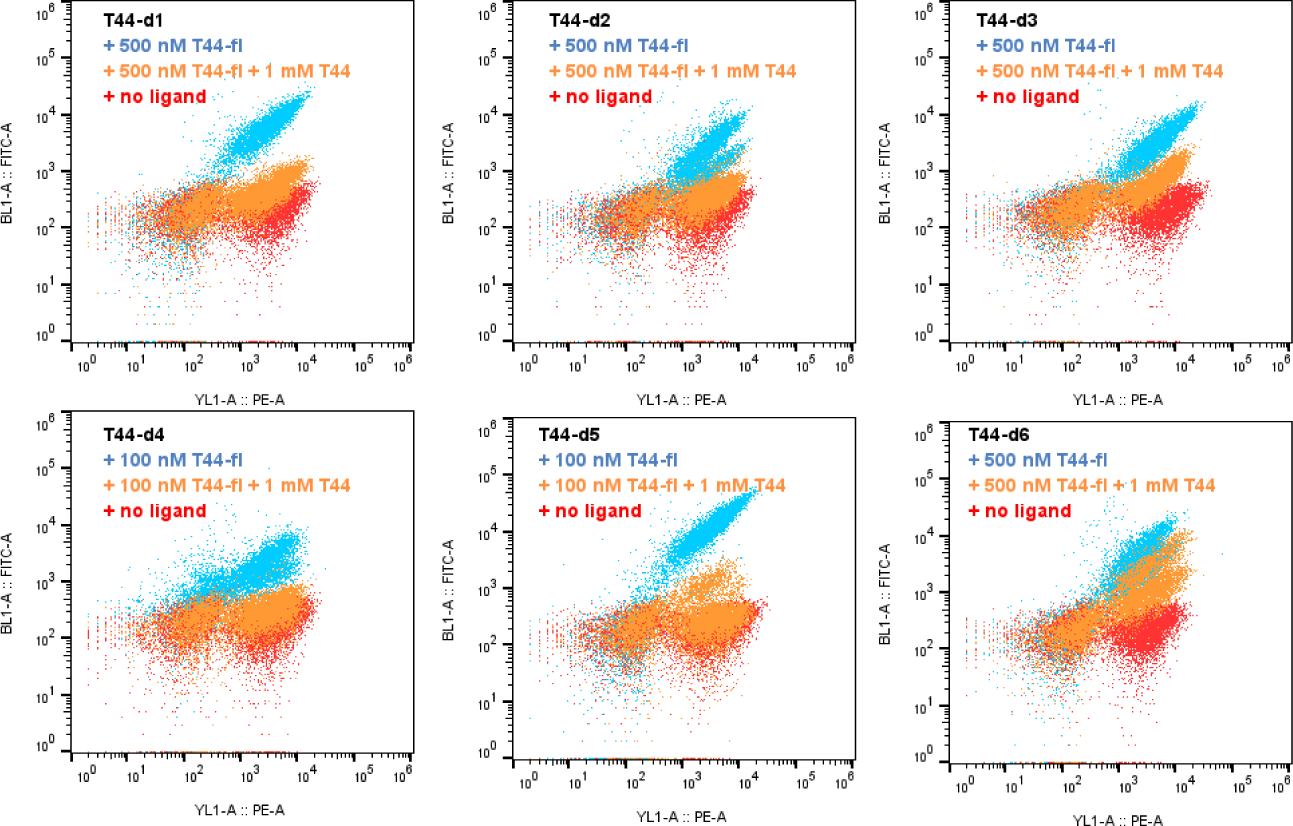
Competition of T44 binding on the yeast surface by free ligand. The designs were displayed on the surface of yeast, using T44-fl for binding and free T44 for competition; specific concenadditionaltrations used are marked on the figure. The binder expressed yeast showed a positive PE signal (YL1-A), and the binding yeast showed a positive FITC signal (BL1-A). All binder expressed yeast showed clear strong double positive signals (blue), while weaker FITC signal upon free ligand competition (orange), and only PE signals when no FITC-labeled ligand presence (red).

**Figure S16.**
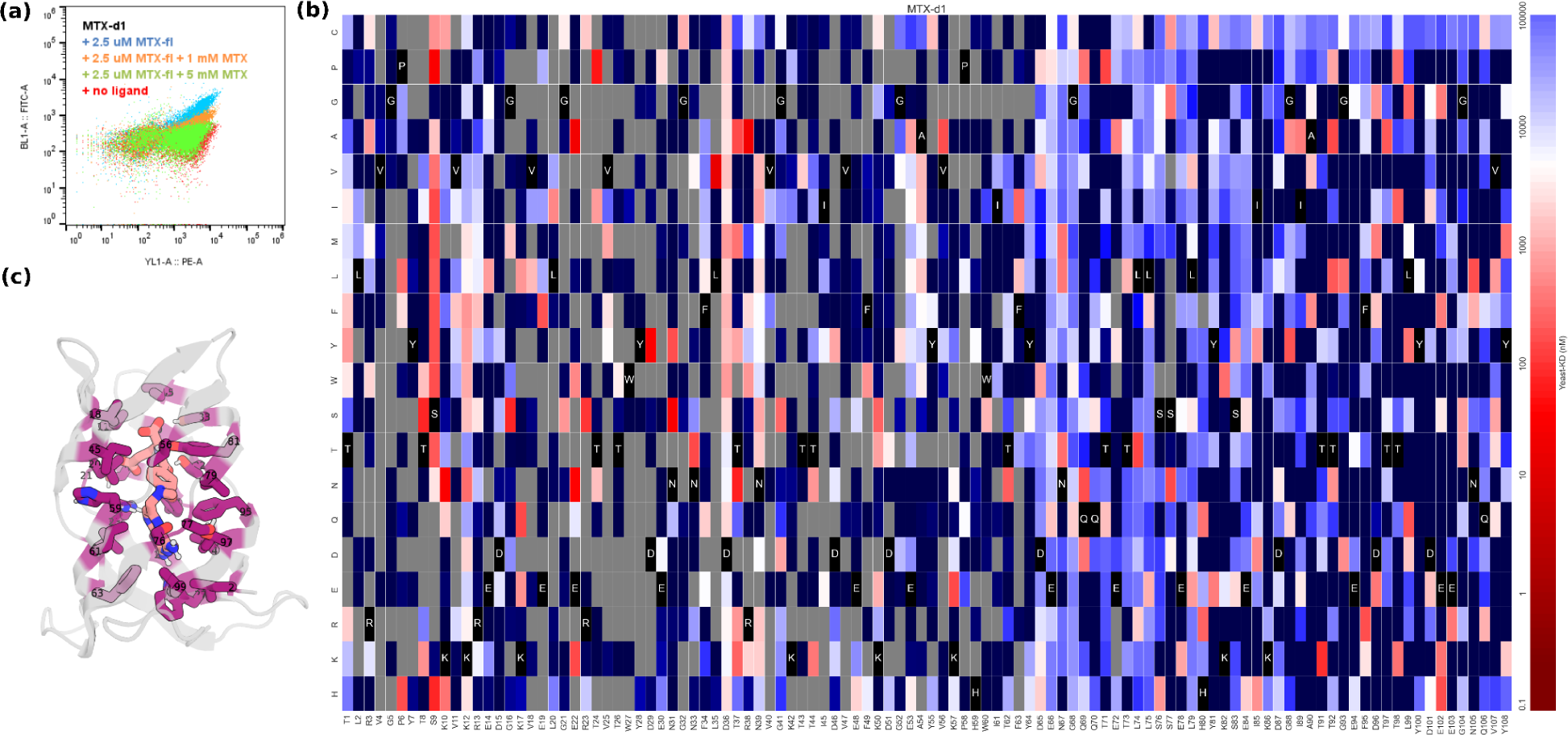
Binding verification of MTX_d1 by site saturation mutagenesis. MTX-d1 showed clear binding competed by free MTX **(a)** on yeast. SSM analysis of MTX supported the designed binding mode **(b-c)**. In the SSM matrix figure **(b)**, the native residues are listed on the x axial, and colored in black. The estimated binding affinity of each mutant is colored based on y axial; the residue choices which may improve binding are more red, while the residues that jeopardize the binding are more blue. For the sorting competition assay in **(a)**, the binder expressed yeast showed a positive PE signal (YL1-A), and the binding yeast showed a positive FITC signal (BL1-A). The binder expressed yeast clearly showed a strong double positive signal (blue), while weaker FITC signal upon free ligand competition (orange and green), and a PE-only signal when no FITC-labeled ligand presence (red). Based on the SSM matrix (**b**), conservative pocket residues are colored in magenta in (**c**).

**Figure S17.**
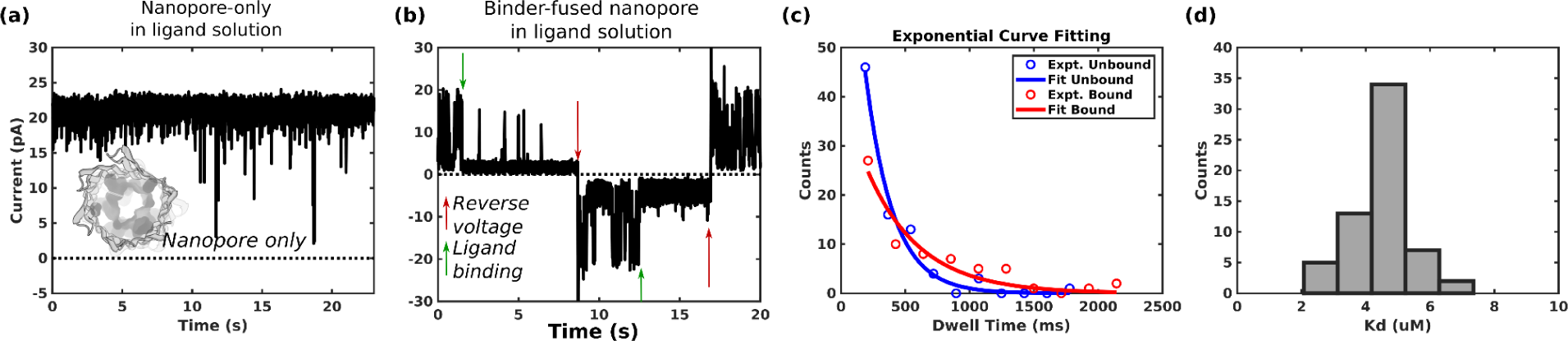
Characterization of binder fused nanopore. The original nanopore did not show obvious blocking in presence of the ligand **(a)**. The binder fused nanopore can transition to off states and can be reversed by reversing the voltage **(b)**. Based on the ligand blocking dwell times (**c**, cutoff of > 100 ms shown here), the Kd of the binder-fused nanopore was estimated, which is in good agreement with reported Kd (CHD-r1, 5.3 ± 0.1 μM) **(d)**. See ‘***Estimation of dissociation constant (Kd) from nanopore conductance measurements****’* in Methods).

**Figure S18.**
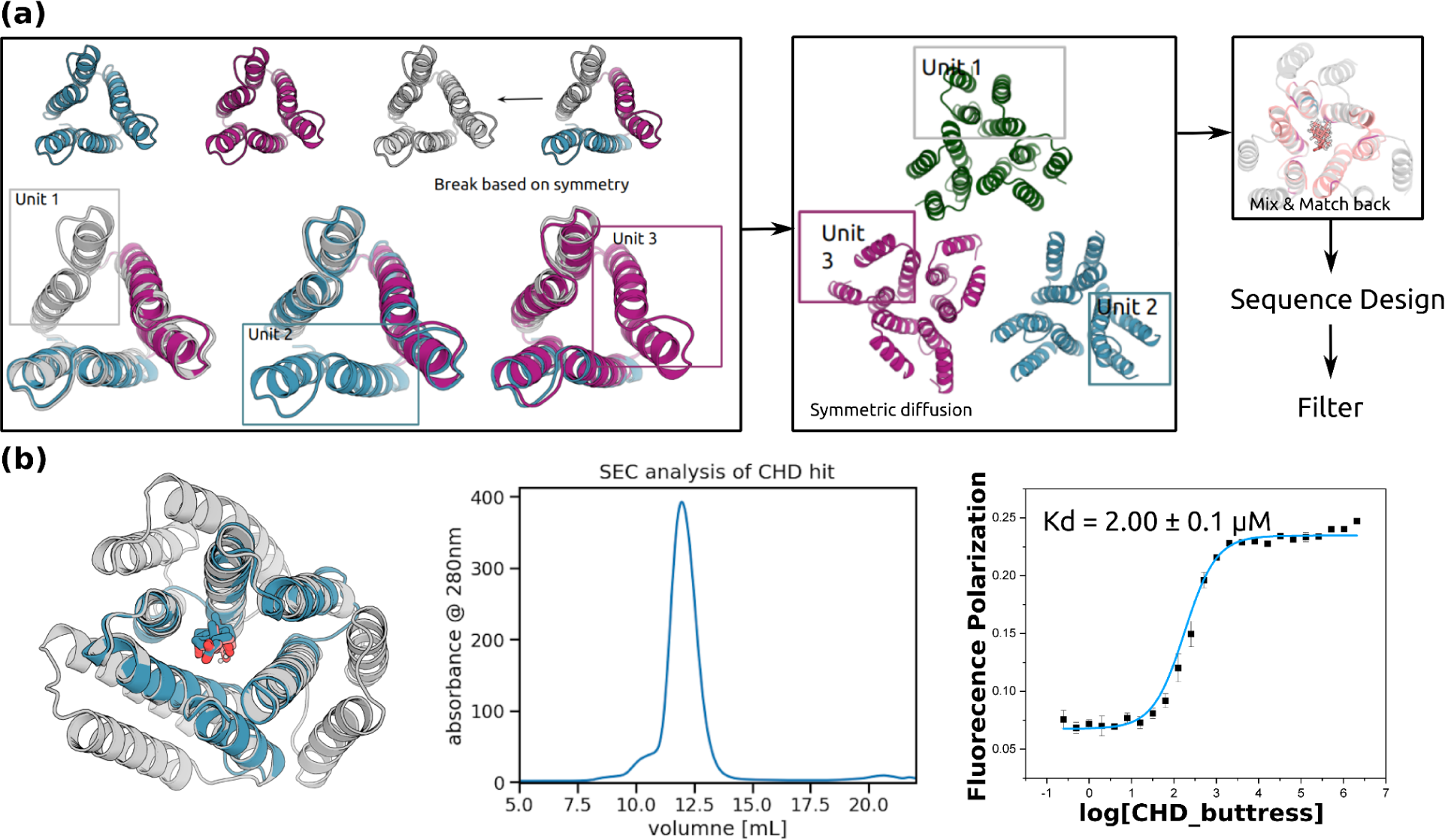
The design procedure for buttrased binder. (a) Design strategy. Single ring designs are broken into repeat units based on the structural symmetry, and two helix buttresses were generated around each repeat protein using symmetric diffusion. The resulting four helical repeat units were pieced back together to generate the final buttrased scaffolds. ProteinMPNN was used to generate sequences. **(b)** The buttressed binder purified as a monodisperse peak and had tighter binding affinity than the original binder, (CHD-r1, 5.3 ± 0.1 μM). Also see **Fig S7** for the co-crystal structure of CHD_buttress with ligand.

**Table S1.** Computational statistics. The numbers of the rotamer used, docks for ligandMPNN or Rosetta design, and final ordered designs from both rounds, and the potential binder, the selected verified binders were reported. ‘Binders selected and verified using purified protein’ meaning, based on predicted affinity and the resource, we choose a smaller number of binders to prioritize from all the NGS-verified binders to purify and verify their binding affinity.

| items | CHD | MTX | T44 | AMA |
| --- | --- | --- | --- | --- |
| Rotamer number | 112 | 28 | 21 | 1 |
| Docks for long design protocols (1st step) | 141,352 | 105,696 | 103,100 | ~100,000 |
| Ordered design (1st step) | 7,443 | 10,174 | 10,990 | 6,852 |
| Docks for long design protocols (2nd step) | ~100,000 | ~100,000 | 489,306 | 62,211 |
| Ordered design (2nd step) | 14,882 (5,574 from PSSM-based Rosetta design methods, 9,308 from ligandMPNN design method) | 8,506 | 5,936 | 8,058 |
| Binders based on NGS (2nd step) | 231 | 1 | 84 | 2 |
| Binders selected and verified using purified protein (2nd step) | 37 | 1 | 10 | 2 |

**Table S2.** Summary of all reported binder in this paper. The generation, characterization, sequence and structure similarity of each binder were reported here. Sequence similarities were calculated using BLASTP, and structural similarities were generated using Dali server(*35*).

| num<br>ber | Binder | Round | Biochemic<br>al<br>characteriz<br>ation | Design<br>method | Structural<br>similarity<br>to PDB50 | Sequence<br>similarity to<br>Uniprot50 | Sequence<br>similarity to<br>closest parent |
| --- | --- | --- | --- | --- | --- | --- | --- |
| 1 | CHD_r1 | 1 | FP,<br>crystallogra<br>phy, Yeast<br>surface<br>display,<br>SEC | PSSM-based<br>Rosetta | 10% to<br>2L6X | 40%idt with 48%<br>coverage to<br>UniRef50<br>RepID=>UniRef50<br>_A0A843FYG7 | n.a |
| 2 | CHD_r2 | 1 | FP, SEC | Repetitive<br>ligandMPN<br>N | 10% to<br>2L6X | 39%idt with 61%<br>coverage to<br>UniRef50<br>RepID=>UniRef50<br>_N9S0L1 | n.a |
| 3 | T44_r1 | 1 | FP, SEC | PSSM-based<br>Rosetta | 10% to<br>6ait-A | 22%idt with 55%<br>coverage to<br>UniRef50<br>RepID=>UniRef50<br>_A0A8H8XJC6 | n.a |
| 4 | T44_r2 | 1 | FP, SEC | PSSM-based<br>Rosetta | 10% to<br>6ait-A | 33%idt with 44%<br>coverage to<br>UniRef50<br>RepID=>UniRef50<br>_A0A1B1E0J0 | n.a |
| 5 | AMA_r<br>1 | 1 | Yeast<br>surface<br>display | PSSM-based<br>Rosetta | 14% to<br>5igh-A | 37%idt with 49%<br>coverage to<br>UniRef50<br>RepID=>UniRef50<br>_A0A1G5S3S9 | n.a |
| 6 | AMA_r<br>2 | 1 | Yeast<br>surface<br>display | Repetitive<br>ligandMPN<br>N | 12% to<br>6veo-A | 35%idt with 51%<br>coverage to<br>UniRef50<br>RepID=>UniRef50<br>_A0A517Y9K7 | n.a |
| 7 | MTX_r1 | 1 | Yeast<br>surface<br>display | Repetitive<br>ligandMPN<br>N | 4%<br>70kn-A | 36%idt with 59%<br>coverage to<br>UniRef50<br>RepID=>UniRef50<br>_A0A1Y3NG75 | n.a |
| 8 | MTX_r2 | 1 | Yeast<br>surface<br>display | Repetitive<br>ligandMPN<br>N | 9% to<br>70kn-A | 32%idt with 52%<br>coverage to<br>UniRef50<br>RepID=>UniRef50<br>_A0A524MJS0 | n.a |
| 9 | CHD_d1 | 2 | FP, SEC,<br>yeast<br>surface<br>display | Repetitive<br>ligandMPN<br>N | n.a | 26%idt with 40%<br>coverage to<br>UniRef50<br>RepID=>UniRef50<br>_A0A7X1B1Q9 | 30%idt with 54%<br>coverage to<br>CHD_r1 |
| 10 | CHD_d2 | 2 | FP, SEC,<br>yeast<br>surface<br>display | Repetitive<br>ligandMPN<br>N | n.a | 29%idt with 42%<br>coverage to<br>UniRef50 | 28%idt with 55%<br>coverage to<br>CHD_r1 |
|  |  |  |  |  |  | RepID=>UniRef50_A0A7V5LIS7 |  |
| 11 | CHD_d3 | 2 | FP, SEC,<br>yeast<br>surface<br>display | Repetitive<br>ligandMPN<br>N | n.a | 33%idt with 41%<br>coverage to<br>UniRef50<br>RepID=>UniRef50_A0A5N6LWE3 | 35%idt with 65%<br>coverage to<br>CHD_r2 |
| 12 | CHD_d4 | 2 | FP, SEC,<br>yeast<br>surface<br>display | Repetitive<br>ligandMPN<br>N | n.a | 32%idt with 45%<br>coverage to<br>UniRef50<br>RepID=A0A0B1R<br>LI2_9NOCA | 42%idt with 58%<br>coverage to<br>CHD_r1 |
| 13 | CHD_d5 | 2 | FP, SEC,<br>yeast<br>surface<br>display | PSSM-based<br>Rosetta | n.a | 29%idt with 45%<br>coverage to<br>UniRef50<br>RepID=>UniRef50_A0A7W1U1H0 | 44%idt with 67%<br>coverage to<br>CHD_r1 |
| 14 | CHD_d6 | 2 | FP, SEC,<br>yeast<br>surface<br>display | Repetitive<br>ligandMPN<br>N | n.a | 30%idt with 37%<br>coverage to<br>UniRef50<br>RepID=>UniRef50_A0A177KWZ7 | 37%idt with 58%<br>coverage to<br>CHD_r2 |
| 15 | CHD_d7 | 2 | FP, SEC,<br>yeast<br>surface<br>display | Repetitive<br>ligandMPN<br>N | n.a | 26%idt with 51%<br>coverage to<br>UniRef50<br>RepID=>UniRef50_A0A350JEW0 | 26%idt with 41%<br>coverage to<br>CHD_r1 |
| 16 | CHD_d8 | 2 | FP, SEC,<br>yeast<br>surface<br>display | Repetitive<br>ligandMPN<br>N | n.a | 26%idt with 32%<br>coverage to<br>UniRef50<br>RepID=>UniRef50<br>_A0A256Y4E2 | 36%idt with 55%<br>coverage to<br>CHD_r2 |
| 17 | CHD_d9 | 2 | FP, SEC,<br>yeast<br>surface<br>display | Repetitive<br>ligandMPN<br>N | n.a | 28%idt with 42%<br>coverage to<br>UniRef50<br>RepID=>UniRef50<br>_A0A3Q3H3H9 | 40%idt with 72%<br>coverage to<br>CHD_r1 |
| 18 | CHD_d1<br>0 | 2 | FP, SEC,<br>yeast<br>surface<br>display | Repetitive<br>ligandMPN<br>N | n.a | 32%idt with 37%<br>coverage to<br>UniRef50<br>RepID=>UniRef50<br>_A0A5C3FD80 | 36%idt with 79%<br>coverage to<br>CHD_r2 |
| 19 | CHD_d1<br>1 | 2 | FP, SEC,<br>yeast<br>surface<br>display | Repetitive<br>ligandMPN<br>N | n.a | 29%idt with 49%<br>coverage to<br>UniRef50<br>RepID=>UniRef50<br>_A0A843BT52 | 38%idt with 60%<br>coverage to<br>CHD_r1 |
| 20 | CHD_d1<br>2 | 2 | FP, SEC,<br>yeast<br>surface<br>display | Repetitive<br>ligandMPN<br>N | n.a | 28%idt with 41%<br>coverage to<br>UniRef50<br>RepID=>UniRef50<br>_A0A2Z5T4F6 | 51%idt with 86%<br>coverage to<br>CHD_r1 |
| 21 | CHD_d1<br>3 | 2 | FP, SEC,<br>yeast<br>surface<br>display | Repetitive<br>ligandMPN<br>N | n.a | 32%idt with 44%<br>coverage to<br>UniRef50 | 28%idt with 51%<br>coverage to<br>CHD_r1 |
|  |  |  |  |  |  | RepID=>UniRef50_A0A0M2PYL4 |  |
| 22 | CHD_d1<br>4 | 2 | FP, SEC,<br>yeast<br>surface<br>display | Repetitive<br>ligandMPN<br>N | n.a | 31%idt with 33%<br>coverage to<br>UniRef50<br>RepID=>UniRef50_A0A1I2G9I0 | 36%idt with 64%<br>coverage to<br>CHD_r2 |
| 23 | CHD_d1<br>5 | 2 | FP, SEC,<br>yeast<br>surface<br>display | Repetitive<br>ligandMPN<br>N | n.a | 42%idt with 64%<br>coverage to<br>UniRef50<br>RepID=>UniRef50_A0A6P8C3U5 | 40%idt with 66%<br>coverage to<br>CHD_r2 |
| 24 | CHD_d1<br>6 | 2 | FP, SEC | Repetitive<br>ligandMPN<br>N | n.a | 23%idt with 44%<br>coverage to<br>UniRef50<br>RepID=>UniRef50_I1C0H3 | 29%idt with 56%<br>coverage to<br>CHD_r1 |
| 25 | CHD_d1<br>7 | 2 | FP, SEC | Repetitive<br>ligandMPN<br>N | n.a | 28%idt with 50%<br>coverage to<br>UniRef50<br>RepID=>UniRef50_K1PVQ4 | 33%idt with 67%<br>coverage to<br>CHD_r2 |
| 26 | CHD_d1<br>8 | 2 | FP, SEC | Repetitive<br>ligandMPN<br>N | n.a | 42%idt with 52%<br>coverage to<br>UniRef50<br>RepID=>UniRef50_A0A4Y2LRW9 | 56%idt with 72%<br>coverage to<br>CHD_r1 |
| 27 | CHD_d1<br>9 | 2 | FP, SEC | Repetitive<br>ligandMPN<br>N | n.a | 26%idt with 44%<br>coverage to<br>UniRef50<br>RepID=>UniRef50<br>_A0A3Q3H3H9 | 35%idt with 50%<br>coverage to<br>CHD_r1 |
| 28 | CHD_d2<br>0 | 2 | FP, SEC | PSSM-based<br>Rosetta | n.a | 34%idt with 64%<br>coverage to<br>UniRef50<br>RepID=>UniRef50<br>_A0A654GKB2 | 56%idt with 78%<br>coverage to<br>CHD_r2 |
| 29 | CHD_d2<br>1 | 2 | FP, SEC | Repetitive<br>ligandMPN<br>N | n.a | 24%idt with 55%<br>coverage to<br>UniRef50<br>RepID=>UniRef50<br>_A0A7R8ZID3 | 53%idt with 78%<br>coverage to<br>CHD_r1 |
| 30 | CHD_d2<br>2 | 2 | FP, SEC | Repetitive<br>ligandMPN<br>N | n.a | 31%idt with 45%<br>coverage to<br>UniRef50<br>RepID=>UniRef50<br>_A0A173XYP3 | 32%idt with 45%<br>coverage to<br>CHD_r1 |
| 31 | CHD_d2<br>3 | 2 | FP, SEC | PSSM-based<br>Rosetta | n.a | 22%idt with 28%<br>coverage to<br>UniRef50<br>RepID=>UniRef50<br>_A0A806DV92 | 33%idt with 55%<br>coverage to<br>CHD_r1 |
| 32 | CHD_d2<br>4 | 2 | FP, SEC | Repetitive<br>ligandMPN<br>N | n.a | 27%idt with 35%<br>coverage to<br>UniRef50 | 39%idt with 61%<br>coverage to<br>CHD_r1 |
|  |  |  |  |  |  | RepID=>UniRef50<br>_UPI000E68460F |  |
| 33 | CHD_d2<br>5 | 2 | FP, SEC | PSSM-based<br>Rosetta | n.a | 30%idt with 36%<br>coverage to<br>UniRef50<br>RepID=>UniRef50<br>_A0A6G1GIV4 | 29%idt with 56%<br>coverage to<br>CHD_r2 |
| 34 | CHD_d2<br>6 | 2 | FP, SEC | Repetitive<br>ligandMPN<br>N | n.a | 41%idt with 57%<br>coverage to<br>UniRef50<br>RepID=>UniRef50<br>_UPI0016673335 | 37%idt with 62%<br>coverage to<br>CHD_r1 |
| 35 | CHD_d2<br>7 | 2 | FP, SEC | PSSM-based<br>Rosetta | n.a | 34%idt with 46%<br>coverage to<br>UniRef50<br>RepID=>UniRef50<br>_A0A3D4MUW3 | 36%idt with 57%<br>coverage to<br>CHD_r2 |
| 36 | CHD_d2<br>8 | 2 | FP, SEC | Repetitive<br>ligandMPN<br>N | n.a | 28%idt with 32%<br>coverage to<br>UniRef50<br>RepID=>UniRef50<br>_A0A7X7GV99 | 37%idt with 60%<br>coverage to<br>CHD_r2 |
| 37 | CHD_d2<br>9 | 2 | FP, SEC | PSSM-based<br>Rosetta | n.a | 28%idt with 36%<br>coverage to<br>UniRef50<br>RepID=>UniRef50<br>_A0A6S7ITZ3 | 39%idt with 61%<br>coverage to<br>CHD_r1 |
| 38 | CHD_d3<br>0 | 2 | FP, SEC | PSSM-based<br>Rosetta | n.a | 27%idt with 39%<br>coverage to<br>UniRef50<br>RepID=>UniRef50<br>_A0A518K765 | 48%idt with 64%<br>coverage to<br>CHD_r1 |
| 39 | CHD_d3<br>1 | 2 | FP, SEC | PSSM-based<br>Rosetta | n.a | 34%idt with 32%<br>coverage to<br>UniRef50<br>RepID=>UniRef50<br>_A0A3M7S8C3 | 35%idt with 51%<br>coverage to<br>CHD_r2 |
| 40 | CHD_d3<br>2 | 2 | FP, SEC | Repetitive<br>ligandMPN<br>N | n.a | 34%idt with 45%<br>coverage to<br>UniRef50<br>RepID=>UniRef50<br>_UPI000711C2C5 | 52%idt with 76%<br>coverage to<br>CHD_r2 |
| 41 | CHD_d3<br>3 | 2 | FP, SEC | Repetitive<br>ligandMPN<br>N | n.a | 30%idt with 60%<br>coverage to<br>UniRef50<br>RepID=>UniRef50<br>_A0A5C5V838 | 47%idt with 72%<br>coverage to<br>CHD_r1 |
| 42 | CHD_d3<br>4 | 2 | FP, SEC | PSSM-based<br>Rosetta | n.a | 24%idt with 53%<br>coverage to<br>UniRef50<br>RepID=>UniRef50<br>_A0A4U0VE94 | 54%idt with 77%<br>coverage to<br>CHD_r1 |
| 43 | CHD_d3<br>5 | 2 | FP, SEC | Repetitive<br>ligandMPN<br>N | n.a | 37%idt with 32%<br>coverage to<br>UniRef50 | 41%idt with 63%<br>coverage to<br>CHD_r1 |
|  |  |  |  |  |  | RepID=>UniRef50_A0A1F9HRL8 |  |
| 44 | CHD_d3<br>6 | 2 | FP, SEC | Repetitive<br>ligandMPN<br>N | n.a | 36%idt with 47%<br>coverage to<br>UniRef50<br>RepID=>UniRef50_Q0UZJ0 | 38%idt with 66%<br>coverage to<br>CHD_r1 |
| 45 | CHD_d3<br>7 | 2 | FP, SEC | Repetitive<br>ligandMPN<br>N | n.a | 32%idt with 36%<br>coverage to<br>UniRef50<br>RepID=>UniRef50_A0A7S3WRZ2 | 33%idt with 61%<br>coverage to<br>CHD_r1 |
| 46 | MTX_d<br>1 | 2 | FP, SEC,<br>yeast<br>surface<br>display | Repetitive<br>ligandMPN<br>N | n.a | 32%idt with 36%<br>coverage to<br>UniRef50<br>RepID=>UniRef50_E3HCG3 | 43%idt with 57%<br>coverage to<br>MTX_r1 |
| 47 | AMA_d<br>1 | 2 | SPR, SEC,<br>yeast<br>surface<br>display | PSSM-based<br>Rosetta | n.a | 37%idt with 49%<br>coverage to<br>UniRef50<br>RepID=>UniRef50_A0A1G5S3S9 | 44%idt with 67%<br>coverage to<br>MTX_r1 |
| 48 | AMA_d<br>2 | 2 | SPR, SEC,<br>yeast<br>surface<br>display | Repetitive<br>ligandMPN<br>N | n.a | 35%idt with 51%<br>coverage to<br>UniRef50<br>RepID=>UniRef50_A0A517Y9K7 | 33%idt with 61%<br>coverage to<br>AMA_r2 |
| 49 | T44_d1 | 2 | FP, SEC,<br>yeast<br>surface<br>display | PSSM-based<br>Rosetta | n.a | 34%idt with 43%<br>coverage to<br>UniRef50<br>RepID=>UniRef50<br>_A0A4P7A274 | 67%idt with 83%<br>coverage to<br>T44_r1 |
| 50 | T44_d2 | 2 | FP, SEC,<br>yeast<br>surface<br>display | PSSM-based<br>Rosetta | n.a | 24%idt with 31%<br>coverage to<br>UniRef50<br>RepID=>UniRef50<br>_A0A8H8JH58 | 65%idt with 85%<br>coverage to<br>T44_r1 |
| 51 | T44_d3 | 2 | FP, SEC,<br>yeast<br>surface<br>display | PSSM-based<br>Rosetta | n.a | 28%idt with 25%<br>coverage to<br>UniRef50<br>RepID=>UniRef50<br>_R7RX94 | 68%idt with 84%<br>coverage to<br>T44_r1 |
| 52 | T44_d4 | 2 | FP, SEC,<br>yeast<br>surface<br>display | Repetitive<br>ligandMPN<br>N | n.a | 31%idt with 42%<br>coverage to<br>UniRef50<br>RepID=>UniRef50<br>_A0A7C2MB67 | 54%idt with 74%<br>coverage to<br>T44_r2 |
| 53 | T44_d5 | 2 | FP, SEC,<br>yeast<br>surface<br>display | Repetitive<br>ligandMPN<br>N | n.a | 33%idt with 57%<br>coverage to<br>UniRef50<br>RepID=>UniRef50<br>_A0A7S3ML05 | 48%idt with 65%<br>coverage to<br>T44_r2 |
| 54 | T44_d6 | 2 | FP, SEC,<br>yeast<br>surface<br>display | Repetitive<br>ligandMPN<br>N | n.a | 21%idt with 49%<br>coverage to<br>UniRef50 | 48%idt with 71%<br>coverage to<br>T44_r1 |
|  |  |  |  |  |  | RepID=>UniRef50_A0A6A4XWT8 |  |
| 55 | T44_d7 | 2 | FP, SEC,<br>yeast<br>surface<br>display | Repetitive<br>ligandMPN<br>N | n.a | 31%idt with 37%<br>coverage to<br>UniRef50<br>RepID=>UniRef50_A0A6N6S117 | 69%idt with 88%<br>coverage to<br>T44_r1 |
| 56 | T44_d8 | 2 | FP, SEC,<br>yeast<br>surface<br>display | Repetitive<br>ligandMPN<br>N | n.a | 31%idt with 35%<br>coverage to<br>UniRef50<br>RepID=>UniRef50_A0A6I7PU97 | 52%idt with 73%<br>coverage to<br>T44_r2 |
| 57 | T44_d9 | 2 | FP, SEC,<br>yeast<br>surface<br>display | Repetitive<br>ligandMPN<br>N | n.a | 30%idt with 40%<br>coverage to<br>UniRef50<br>RepID=>UniRef50_A0A397TFF8 | 52%idt with 75%<br>coverage to<br>T44_r2 |
| 58 | T44_d10 | 2 | FP, SEC,<br>yeast<br>surface<br>display | Repetitive<br>ligandMPN<br>N | n.a | 26%idt with 34%<br>coverage to<br>UniRef50<br>RepID=>UniRef50_A0A1H8UAJ6 | 46%idt with 70%<br>coverage to<br>T44_r1 |

\*Structural similarity described as %idt

**Table S3.**
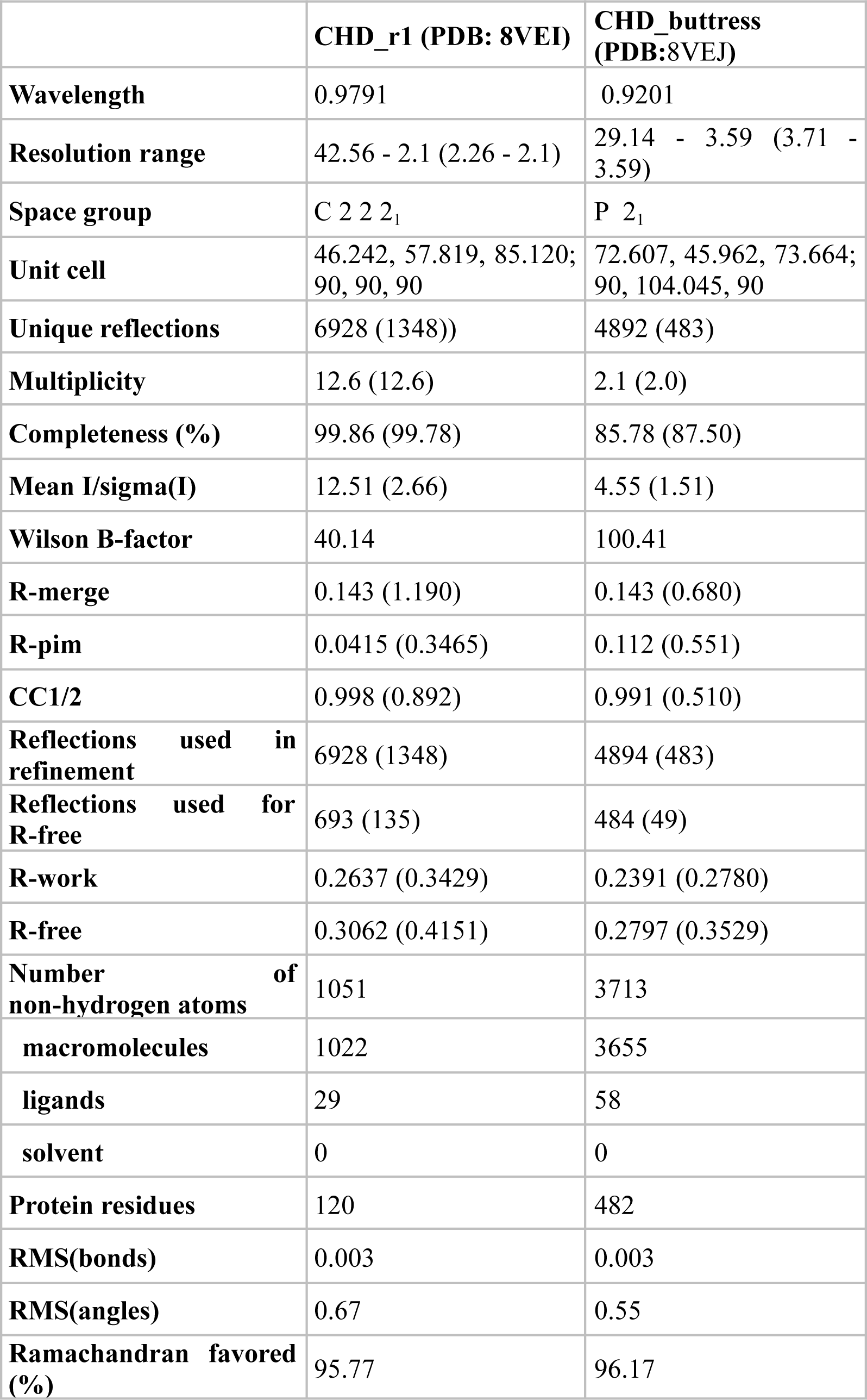
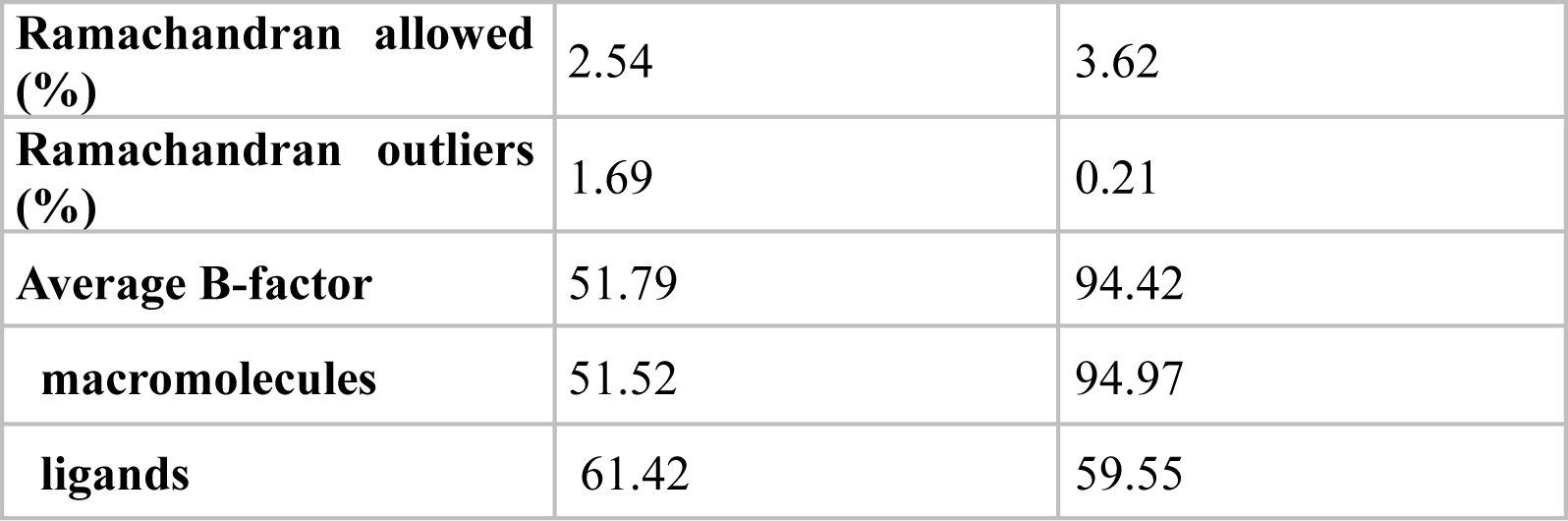
Data collection and refinement statistics for co-crystal structures.

**Table S4.**
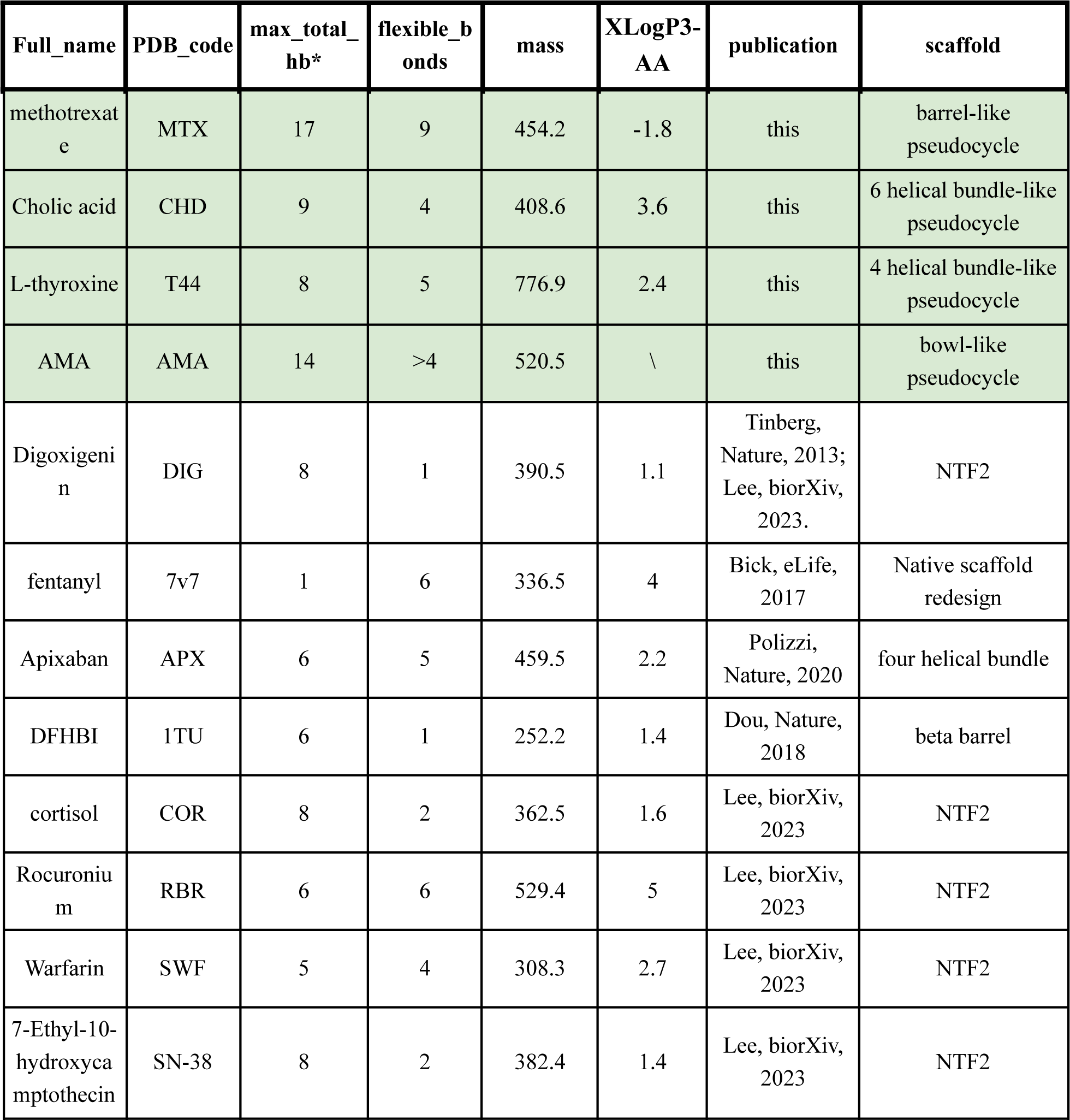
Ligand complexity characterization. The features of Ligands from previous and current work (green) of SM binder designs are extracted from PubChem(*36*) or counted (for AMA) and listed here. Only ligands with binders with affinity reported and some binding pose validations (SSM, competition, or crystallization) are included here. *max_total_hb equals addition of the number of hydrogen bond donors and the number of hydrogen bond acceptors.

| Full_name | PDB_code | max_total_hb* | flexible_bonds | mass | XLogP3-AA | publication | scaffold |
| --- | --- | --- | --- | --- | --- | --- | --- |
| methotrexate | MTX | 17 | 9 | 454.2 | -1.8 | this | barrel-like pseudocycle |
| Cholic acid | CHD | 9 | 4 | 408.6 | 3.6 | this | 6 helical bundle-like pseudocycle |
| L-thyroxine | T44 | 8 | 5 | 776.9 | 2.4 | this | 4 helical bundle-like pseudocycle |
| AMA | AMA | 14 | >4 | 520.5 | \ | this | bowl-like pseudocycle |
| Digoxigenin | DIG | 8 | 1 | 390.5 | 1.1 | Tinberg, Nature, 2013; Lee, biorXiv, 2023. | NTF2 |
| fentanyl | 7v7 | 1 | 6 | 336.5 | 4 | Bick, eLife, 2017 | Native scaffold redesign |
| Apixaban | APX | 6 | 5 | 459.5 | 2.2 | Polizzi, Nature, 2020 | four helical bundle |
| DFHBI | 1TU | 6 | 1 | 252.2 | 1.4 | Dou, Nature, 2018 | beta barrel |
| cortisol | COR | 8 | 2 | 362.5 | 1.6 | Lee, biorXiv, 2023 | NTF2 |
| Rocuronium | RBR | 6 | 6 | 529.4 | 5 | Lee, biorXiv, 2023 | NTF2 |
| Warfarin | SWF | 5 | 4 | 308.3 | 2.7 | Lee, biorXiv, 2023 | NTF2 |
| 7-Ethyl-10-hydroxycamptothecin | SN-38 | 8 | 2 | 382.4 | 1.4 | Lee, biorXiv, 2023 | NTF2 |

## Supplementary Materials

### Materials and Methods

#### Stepwise SM binder design pipeline in detail

All scripts on stepwise SC-optimizing SM binder design pipeline are available at github (repository will be public after manuscript acceptance): https://github.com/LAnAlchemist/Pseudocycle_small_molecule_binder.

All binders were designed using the same computational pipeline unless specified otherwise.

#### Ligand preparation for computational processing

The 3D structure of the ligands were first taken from PDB or generated using ChemDraw 3D. The hydrogens were added using OpenBabel(*37*), and edited with VMD(*38*) for correct chemical structures and protonation states. Around 300-700 rotamers were generated for each ligand using RDKit(*23*). All rotamers were then relaxed using Rosetta with constraint, and scored for ddG. The scored rotamers were also aligned using RDKit, function ‘Chem.rdMolAlign’, and clustered based on r.m.s.d, using cutoff at 1.5 Å. The rotamer with lowest Rosetta energy from each cluster was selected as the starting point. The final selected rotamers are all physically allowed, but not necessarily the lowest-energy rotamers among all possible rotamers. The parameter file of each ligand based on ‘ref2015’, ‘genpot’(*39*) score functions were generated using Rosetta. The only exception is AMA, where its single designed structure of AMA was used as the binding confirmation.

#### First step SC-optimizing sampling

The previously published 9838 pseucocles were reduced to 9,703 based on protein length; only proteins with length shorter than 155 aa were used for the following design work due to the limitation of the oligo library synthesis availability. The pocket residues of pseudocycles were annotated using in-house python script which identifies the largest internal cavity bounded by the protein after converting the protein to polyalanine and then identifies all side chain residues contacting this internal cavity.(*11*)

All rotamers of all four ligands were first docked to 9,703 pseudocycles using RIFgen/RIFdock suite(*3*) using the same docking protocol as the previous publication(*11*). Considering the aim of the first sampling step is to identify suitable scaffold for each ligand, to save resources, only 10 docks were requested between each ligand rotamer to each pseudocycle scaffold.

#### Predictor to select best docks

All docks obtained from the previous step were collected and the interfaces (a.k.a ligand and the pocket residues) were designed using a quick design and score Rosetta protocol (‘FastPredictor_v2_talaris_ligand.xml’). The interface except for the previously seeded rifres was quickly packed with big hydrophobic residues (a.k.a Val, Leu, ILE, PHE, TYR) using score function ‘talaris2014’. No Rosetta relaxation was performed, and only ‘contact_molecular_surface’ (CMS) was scored to save time. On average, 10 docks takes ∼ 1 CPU second to score. The CMS scores were collected and ranked from highest to lowest. Docks at different CMS ranks were pulled out for manual inspection, and a cutoff CMS score was selected based on how well the ligand contacts with the scaffold. Usually 30-50% of the docks were dropped at this stage.

The selected docks were designed and scored again using another relatively quick design and score Rosetta protocol. The interface except for the previously seeded rif res were packed using layer design protocol (‘LA_quick_design_select_dock_genpot.xml’). On average, 1 dock takes ∼

10 CPU seconds to score. ‘CMS’, ‘dsasa’, ‘ddg’ (a.k.a ‘ddg_norepack’ in the protocol), ‘holes_around_lig’ were used as major criterias to select final docks for design. The features were scored and ranked, and cutoffs were selected by manual inspection of docks at different ranks. The low cutoff, low cutoff, high cutoff, and high cutoff were selected for the abovementioned metrics, respectively.

All docks post first predictor were also taken and measured the potential ligand tail atom burial to avoid designs with too buried ligands.

All scores generated from predictor protocol scripts were found to be statistically highly correlated with the scores generated from the actual design protocols, which indicates the 2-step predictor is a good method to rank docks and select good docks to improve computational design efficiency (Fig S5).

#### PSSM-based Rosetta design protocol

One PSSM file was generated for each pseudocycle scaffold using ProteinMPNN(*13*). 100 sequences were generated for each pseudocycle scaffold using ProteinMPNN with temperature at 0.2 to increase sequence diversity. The PSSM score at each position of each amino acid type was calculated through comparing proteinMPNN generated frequency (p_observed) to that of BLOSUM62 amino acid background frequency (p_background). The calculation equation is:

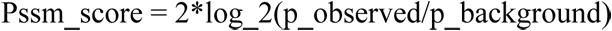

Where p_observed is the frequency of the particular amino acid type observed at the specified location.

Two rounds of FastDesign were performed with the sequence biased using PSSM (‘design_ligand_full_noHBNet_1.py’). For a 120-aa protein, usually it takes 15 min CPU time for each design and score. Because the quality of both the protein model (as assessed by AlphaFold2(*14*), AF2), and the ligand docks (selected by the two-step ‘predictor’) were high at this point, for both protocols, strong constraints were added to disallow significant movements of the protein backbone or the ligand docking conformation.

#### Iterative ligandMPNN design protocol

The output of the PSSM-based Rosetta designed proteins were used as input for repetitive ligandMPNN design protocol (‘ligMPNN_FR_silent_in.py’). All short range interactions seeded from the previous design and RIFdock (polar interactions, π-π interactions) were kept in the first round of ligandMPNN design, while all residues were allowed for design during the second and third round of ligandMPNN design. Rosetta minimization with strong ligand-protein constraints were performed between ligandMPNN runs to remove potential unideal local interactions while preserving the ligand docking pose and protein backbone conformations. Temperature of 0.2 and polar/nonpolar bias were used to increase the sequence diversity and polar interface building bias.

#### Selection of designs for ordering

All designs were pooled together and filtered with the same criteria. Rosetta metrics including ‘CMS’, ‘ddg2’, ‘hole_around_lig’, ‘dsasa’, ‘total_hb_to_lig’ were used for design judgements. Designs with different scores of above-mentioned metrics were manually inspected for low cutoff, high cutoff, high cutoff, low cutoff, and low cutoff selection, respectively. The proteins of designs passed Rosetta filters were then first scored using AF2, and only designs with pLDDT (predicted local distance difference test) above 85, predicted C7-r.m.s.d smaller than 1 or 1.5 Å were ordered.

#### The second step SC-optimizing sampling

The pseudocycle scaffold-ligand pair identified from the first stem sampling and showed potential binding in the laboratory (**Fig 1c-f, S2, S3, S6**) were used for the second computational sampling step. The same second step of the sampling pipeline was used for all ligands. First, the hit scaffolds were taken and resampled through generating 1000 sequences using proteinMPNN(*13*) and folded using AF2(*14*). Only AF2 models (model 4) predicted to be within

Cɑ-r.m.s.d 3 Å, plDDT above 90, predicted template modeling score (pTM) above 0.65 were taken as the newly resampled models for next docking and design step.

The same docking procedure was used between each ligand rotamers and their hit scaffolds, except for 1000-1500 docks were requested. The same two-step predictor protocol was used to identify good docks for actual design protocols, which the selection criteria were changed to using the scores from the previous hits. The same sequence design protocols and filter steps were used, except for most of the selection criteria were taken from the previous hits.

Once optimal scaffold topologies were identified for each ligand in the first round, the success rate in the second design round was significantly higher than in previous studies using a single protein family for ligand binding(*3*–*5*, *7*, *8*). For easier targets, such as CHD, after NGS analysis, we confirmed 231 binders from 14,882 designs, with a 1.55% success rate. For the flexible T44, 84 binders were obtained based on NGS data from 5,936 designs, a 1.42% success rate. Both are significantly higher than previous studies where a single family of proteins were used for ligand binder design(*3*, *6*, *8*).

With further data on optimal pseudocycle geometries for different target ligands, it should become possible to identify optimal scaffolds for a given ligand computationally which could obviate the need for the initial design round, considerably streamlining the design process (the ‘contact_molecular_surface’, measuring protein-ligand contacts, is significantly higher for the round 1 designs with binding activity, but more data is needed to set thresholds; Fig S1c).

#### Preparation of the FITC-labeled SM target

Free CHD (1133503-200MG), T44 (T2376-1G), MTX (454126-100MG) were purchased from Sigma-Aldrich (St. Louis, MO). MTX-fl was purchased from Thermo Fisher (M1198MP, Bothwell, WA). MTX-btn (M260670) was purchased from Toronto Research Chemicals. CHD-fl was purchased from Life Sciences (#451041). Biotinylated AMA were synthesized, purified, and verified by WuXi Chemistry AppTec, Inc.

Preparation of the FITC-labeled T44 was performed as previously described with some slight modifications(*40*). Briefly, L-thyroxine was incubated directly with N-hydroxysuccinimide-fluorescein (2 eq.) and N,N-diisopropoylethylamine (4 eq.) in dimethylformamide (DMF) for 4 hours. Following conversion to the fluorescent product, which was confirmed by mass spectrometry, the reaction was rotavapored to remove DMF, resuspended in acetonitrile and water, and purified by reverse-phase high-performance liquid chromatography to yield FITC-labeled T44.

All ligands were stored following commercial instructions, or in 30% (v/v) dimethyl sulfoxide (DMSO)/water at ™20 ℃ for long-term storage.

#### High-throughput methods for binder identification

All library oligos were purchased from Twist Bioscience (South San Francisco, CA). The KAPA HiFi HotStart Uracil+ReadyMix (KK) and KAPA HiFi HotStart Kit (KK2101) were purchased from Roche (Basel, Switzerland). All gene fragments were purchased from Twist Bioscience (South San Francisco, CA) or IDT (Coralville, IA). The EvaGreen Dye, 20 x in water (#31000), was purchased from Biotium (Fremont, CA). The USER enzyme (NEB#M5508) and the NEBNext End Repair Module (E6050L) were purchased from New England BioLabs (Ipswich, MA). All DNA purification kits were purchased from QIAgen (Hilden, Germany). All chemicals and consumables were purchased from Thermo Fisher(Bothell, WA), unless specified otherwise. The anti-cMyc-R,Phycoerythrin (anti-cMyc-PE) was purchased from Cell Signaling Technologies (Danvers, MA). All tagged and free ligands were described in ‘**Preparation of the FITC-labeled SM target**’ in Methods.

#### Yeast surface display library preparation

The designed proteins were reverse transcribed using DNAworks(*41*) with avoidance of common restriction enzyme cut sites. The nucleotide sequences were then broken into two parts (5’ part, 3’ part) with common complementary sequences in the middle for assembly using an in-house python script. The designs were separated into different groups of around 1000 based on length to avoid quantitative PCR (qPCR) bias caused by oligo length differences. The primers complementary to the pETCON3 yeast surface display vectors were added to the 5’ of the 5’ part and 3’ of the 3’ part. Inner primers with AA, TT in the middle were padded to the 3’ of the 5’ part and 5’ of the 3’ part to facilitate library PCR. The 250 and 300 nt oligo pools were ordered from Twist Bioscience depending on the protein length.

The purchased oligo pools were dissolved in water to prepare 100 ng/μL stock solution and stored at ™20 ℃. The oligo pool stock was diluted to 2.5 ng/μL in water for qPCR experiments. The individual 5’-parts and 3’-parts were amplified using designed primers following commercially available protocol from KAPA HiFi HotStart Uracil+ReadyMix. DNA was purified using a PCR purification kit in between the qPCR runs. The inner primers of individual parts were removed using USER enzymes and the NEBNext End Repair Module following commercially available protocols. After DNA purification, the digested 5’-parts and 3’-parts were assembled using the same qPCR protocol, and the assembled oligo pools were checked using DNA electrophoresis. The designed DNA bands were cut from gels and purified for amplification. Roughly 4-6 μg of oligo pools were used for electrocompetent yeast transformation for a library with complexity between 5,000-10,000. The oligo pool and the linearized pETCON3 vector were added to yeast at a ratio of 3:1 and the electro-transfer protocol for the yeast library was used for transformation following the previously published protocol(*42*).

#### Yeast surface display experiment for enrichment of the potential binders

The previously established yeast surface display protocol was followed with minor modification(*42*). PBSF, PBS with 0.1% (w/v) Bovine Serum Albumin (BSA), were used as all yeast sorting buffers. EBY1 were used for all yeast surface display experiments. All library fluorescence-activated cell sorting (FACS) experiments were performed on a Sony SH800S Cell Sorter using 100-μm chip, which is equipped with laser 488 nm and 561 nm.

For all yeast surface display libraries, one round of expression sort was first performed to enrich the binders with acceptable expression level. The binder-expressing yeast were stained at 1:50 (v/v) ratio with anti-cMyc-FITC for 30 min at 4 ℃ for expression sort; yeast with increased FITC signal, indicating expression, were collected for further binding assays. After expression sort, the collected yeast were grown up in C-Trp-Ura-2% glucose (CTUG) medium again, and expressed overnight at 30 ℃ in a shaker in SGCAA medium for binder expression.

If the ligands were tagged with FITC, the expressed yeast was first stained with 1:35 (v/v) anti-cMyc-PE for 45 min at 4 °C, followed by two washes using PBSF. Then the stained yeast was incubated with FITC-labeled ligands for 60-120 min at 4 °C. After two more washes, the stained yeast was sorted. The expressed-only yeast was used as negative control, while the yeast population with positive signals from both FITC and PE channels above the negative control samples were collected and grown up in CTUG medium. After 1-2 days of growth, the collected yeasts were expressed again in SGCAA medium and sorted again for binder enrichment through a similar above-described method. Usually it takes two to three sorts to have clear enrichment of the binding populations. If the binding population was strong (above 2%), a titration may be performed to select the strongest binders. Titration sorts were performed for CHD and T44 for both round1 and round2 binding assays. Usually 5 μM FITC-labeled ligands were used for sort1. The concentration would be dropped according to the positive signal intensity at sort2 and sort3. Usually 100 pM - 5 uM of FITC-labeled ligands were used for titration sorts.

If the ligands were tagged with biotin, the expressed yeast was always stained with anti-cMyc-FITC at 1:50 (v/v) for 35 min at 4 °C. The first and second sort were usually performed with avidity, meaning, 1 μM of biotinylated ligands were added to yeast with 0.25 μM anti-Streptavidin-R,Phycoerythrin (SAPE) at the same time for 120 min at 4 °C during the ligand binding step. If the binding population was strong enough, the second, or third round of sorting would be performed without avidity, meaning 100 nM-10 μM of biotinylated ligands were added to design-expressing yeast for incubation for 120 min at 4 °C. After two washes using PBSF, SAPE would be added to stain the ligand-bound yeast at 1:50 (v/v) for 35 min at 4 °C.

Cultures from expression, and all sorting steps were collected for MiSeq analysis. All yeast sorting figures were prepared using FlowJo v10 (FlowJo, Inc).

#### Next-generation sequencing data analysis

All sequences were assembled and matched using an in-house script, and the git repository is at: https://github.com/feldman4/ngs_app.

For sequencing data with titration sorting, the data were analyzed based on previously published scripts to identify potential binders(*43*), and designs with predicted Kd with equal or lower than micromolar range were defined as potential binders, and some of them were selected for individual purification and binding assays, such as SEC, FP, and SPR.

For sequencing data without titration sortings, enrichment factors were calculated based on the reads from sort3 versus the reads from sort2, usually the binders with enrichment factor above 10 and reads from sort3 more than 1000 were ordered individually for binding test.

#### Yeast surface display for individual binder identification

A 120 μg of each gene fragment encoding individual design was mixed with 40 μg of linearized pETCON3 vectors and transferred to chemically competent yeast EBY1 following previously published protocols(*43*). The transformed yeast were first grown in CTUG media in a 96-deep well round bottom plate in plate shaker at 30 °C for 30 hours at 200 g, and expressed in SGCAA media for 14 hours under the same shaking conditions.

The expressed yeast were stained in the same fashion as previously described (see ‘*High-throughput methods for binder identification*’ in Methods) and the binding and expression were checked on an Attune Flow Cytometer equipped with 488 nm and 561 nm lasers. For competition assays, free ligands were added with around 1000 folds higher than the FITC or biotin-labeled ligand at the same time, with the rest of the steps following the same like those described above.

All yeast sorting figures were prepared using FlowJo v10 (FlowJo, Inc).

#### Expression and purification of selected proteins

The major procedures were adapted from a previous publication(*11*). All chemicals and supplies were purchased from Thermo Fisher unless specified otherwise.

The selected designs were reverse-transcribed into DNA using DNAWorks(*41*). Eblocks (IDT or Twist Bioscience) were cloned into a pET29b-derived vector with C-terminal SNAC cleavable His-Tags using commercially available Golden Gate assembly kit (New England Biolabs, Ipswich, MA) and transformed into *Escherichia coli (E. coli)* BL21 strain. All designs were grown in 50 mL autoinduction Terrific Broth media in 250 mL baffled erlenmeyer flasks for production (6 hours at 37 ℃ followed by 24 hours at 18 ℃ shaking at around 200 g in New Brunswick Innova® 44 shakers). Cells for each design culture were harvested using centrifugation and resuspended in 25 mL of lysis buffer (25mM Tris 100 mM NaCl, pH 8, with protease inhibitor tablet) and sonicated to lyse (4.5 minutes sonication, 10 s pulse, 10 s pause, 65% amplitude). After centrifugation for 30 min at 14,000 g, soluble fractions were bound to 1 mL Ni-NTA resin (Qiagen) in a Econo-Pac® gravity column (BIO-RAD) for 1 hour with rotation. The resin was washed with 20 CV (column volume) low salt buffer (50 mM tris, 100 mM NaCl, 10 mM Imidazole, pH 8) and with 20 CV high salt buffer (50 mM tris, 1000 mM NaCl, 10 mM Imidazole, pH 8). Proteins were eluted with 2 CV of elution buffer (20 mM tris, 100 mM NaCl, 500 mM Imidazole, pH 8) and purified on a superdex 75 increase 10/300 GL column connected to ÄKTA protein purification systems in isocratic TBS buffer (20 mM Tris, 100 mM NaCl, pH 8).

For proteins for FP, SPR, and crystallization studies, the Histags were cleaved following on-bead SNAC tag cleavage protocol(*44*). The proteins were prepared using the same way described above until protein binding onto the resin. The column was washed twice with lysis buffer to remove imidazole, followed by equilibrium with 10 CV of SNAC buffer (100 mM CHES, 100 mM acetone oxime, 100 mM NaCl, 500 mM guanidine chloride, pH 8.5). Then the columns were incubated with 20 CV of fresh SNAC cleavage (2 mM NiCl2 in SNAC buffer) overnight at r.t. for SNAC tag on resin cleavage. The resins were then washed with 10 CV of weak elution buffer (20 mM Tris, 100 mM NaCl, 10 mM Imidazole, pH 8). Both SNAC cleavage solution and the weak elution solution were collected, and checked with protein SDS-Page for tag-free proteins. The tag-free proteins were pooled, concentrated, and purified on SEC using the same method described previously.

All purified proteins were verified using mass spectrometry.

#### Fluorescence polarization studies

The protein concentrations were judged using Bradford protein assay (Bio-Rad, Hercules, CA) following commercially available protocols.

All FP studies and analyses were performed following a previously established protocol(*45*) on a Biotek Synergy Neo2 Reader equipped with Dual PMT optical filter cube (part number: 1035108). The proteins were serial diluted and 5 or 20 nM of FITC-labeled ligands were added to the diluted protein. The mixtures were transferred to Corning 384 Well microplate (CLS4515-10EA). After 30 min incubation at 25 ℃ with shaking, the parallel and perpendicular light were read with scale to the highest-intensity well. Two independent FP were performed as replicates.

FP signal was calculated based on below equation:

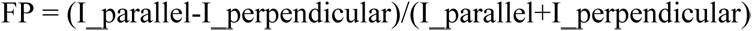

The data were fit using the Origin with GrowthCurve DropWise model to avoid ligand binding saturation.

#### SPR studies

The protein concentrations were judged using Bradford protein assay (Bio-Rad, Hercules, CA) following commercially available instructions. All SPR studies were performed on a Cytiva Biacore 8K SPR system, using a biotin CAPture chip, and in 1x HBS-EP+ (0.1 M HEPES, 1.5 M NaCl, 0.03 M EDTA and 0.5% v/v Surfactant P20, Cytiva) buffer.

All runs were performed using both flow cells, with one reference cell and one experimenting cell.

A start-up run was always first performed with running biotin CAPture reagent (Cytiva) through both cells for 300 seconds contacting, at flow rate of 2 μl/min, following 25 nM biotinylated ligand (biotinylated-AMA) contacting for 180 seconds at at a flow rate of 10 μl/min for flow cell 2. The buffer was then flowed through with 120 seconds contacting and 60 seconds dissociation at a flow rate of 30 μl/min, regeneration solution (3:1 (v/v) 8 M guanidine HCl pH 8, and 10 M NaOH) contacting for 120 seconds at flow rate of 10 μl/min, and lastly washed with buffer.

Blank runs were run before and after analysis runs to set up blanks, the only difference between analysis runs and blank runs was that buffers were used instead of binding proteins.

For analysis runs, the proteins were prepared at 6 concentrations, and tested from low concentration to high concentration sequentially following the same protocol. First, biotin CAPture reagent was flown through both cells for 300 seconds contacting, at a flow rate of 2 μl/min. The 25 nM biotinylated ligand (biotinylated-AMA) contacts for 180 seconds at a flow rate of 10 μl/min for only flow cell 2. Then, the protein solution was flow through for 120 seconds, with 300 seconds disassociation time, at a flow rate of 30 μl/min, to both cells. All 6 different concentrations were flown through sequentially from low to high concentrations. Lastly, the generation solution and wash cycle were performed similar to the start-up round.

The SPR data were analyzed and plotted using Cytiva Biacore Insight Evaluation software.

#### Site-specific mutagenesis SSM studies

SSM studies were performed following the previous protocol(*43*). The SSM library was prepared and sorted following previous protocol (see ‘*Yeast surface display library preparation*’) with minor modification. Each protein residue was mutated to all 20 possible residues, including Cystine. 1 to 5 binders to the same target were pooled together as 1 single SSM library for yeast surface display studies. The only difference during sorting was that the same gate was used for sort 1, 2. Various concentrations of ligands were used for sort2 for titration studies following previously published protocol(*42*). The site-specific Shannon entropy was calculated based on the counts and enrichment following the previously published protocol, and the SSM matrix plots were generated based on the Shannon entropy(*43*).

#### Design of SM binder fused nanopore

The nanopore TMB12_3 from our de novo nanopore design(*46*) was chosen for the design of the cholic acid binder-based nanopore sensor. This nanopore was shown to have a stable conductance at ∼230 pS in a 500 mM NaCl buffer. Also, since the SM binder for cholic acid has 6 helices, a 12 stranded beta-barrel nanopore seemed appropriate for the fusion where 4 strands from the beta-barrel could be fused to 2 helices from the binder leading to a maximum of 6 loop connections between the two proteins. 6 loops with the right lengths would possibly allow for complete blockage of the nanopore opening from a direction perpendicular to the pore and SM binder axis. This is a desirable feature for a binary nanopore sensor where a binding event can be associated with significant current blocking leading to nanopore currents very close to 0 pA. Therefore, the SM binder was placed on one side of the nanopore and roughly aligned to match the closest helical and strand termini that would generate the most optimal fusion without significantly changing the structure of either protein. Complete freedom was allowed for different loop closure solutions based on the orientations of the binder and the nanopore. Further sampling of the binder position was carried out by rotating the binder around the pore axis within an angular range that would allow for different termini distances between the fusion points of the binder and the nanopore. The upper limit on termini distance (between a helix of the binder and a strand of the nanopore) was chosen as 12 Å based on initial sampling which showed that further distances would lead to a different loop closure solution as compared to the starting configuration. Also, the binder was translated along the pore axis within 5-8 Å of the pore opening to allow for different loop conformations upon fusion. The loop closure was carried out using a deep learning model(*25*) for a range of starting termini configurations and their corresponding conformations under different rotations and translations of the binder as described above. Different loop lengths were sampled and final selection for the generated backbones was carried out by filtering out structures with chain breaks. Sequence design was carried out using ProteinMPNN(*13*) and Rosetta total scores were used for final filtering as AF2 was incapable of predicting full structures from single sequences for these types of fusion proteins.

#### Expression and purification of designed binder-based nanopore sensors

Fusion constructs as designed above were ordered as Eblocks from IDT and cloned into a pET29b vector with T7 promoter and kanamycin resistance gene. The proteins were expressed and purified using an auto-induction media as previously described(*47*). All proteins were refolded from inclusion bodies using dodecyl-phosphatidylcholine (DPC) detergent and were subsequently run through a SEC column (Superdex S200 from Cytiva). Appropriate fractions were diluted to nanomolar to picomolar concentrations and tested for membrane insertion and conductance as described below.

#### Conductance measurement in planar lipid bilayers

The conductance measurements were performed as previously described(*47*). All ion-conductance measurements were carried out using the Nanion Orbit 16TC instrument (https://www.nanion.de/products/orbit-16-tc/) on MECA chips. Lipid stock solutions were freshly made in dodecane at a final concentration of 5mg/mL. Di-phytanoyl-phosphatidylcholine (DPhPC) lipids were used for all experiments. Designed proteins were diluted in a buffer containing 0.05% DPC (∼ 1 critical micelle concentration), 25 mM Tris-Cl pH 8.0 and 150 mM NaCl to a final concentration of ∼100 nM. Subsequently, 0.5 μL or less of this stock was added to the cis chamber of the chip containing 200 μL of buffer while simultaneously making lipid bilayers using the in-built rotating stir-bar setup. All measurements were carried out at 25 ℃ and with a positive potential bias of 100 mV. Spontaneous insertions were recorded over multiple rounds of bilayer formation. All chips were washed with multiple rounds of ethanol and water and completely dried before testing subsequent designs. A 500 mM NaCl buffer was used on both sides of the membrane for all current recordings. Raw signals were recorded at a sampling frequency of 5 kHz. Only current recordings from bilayers whose capacitances were in the range 15-25 pF were used for subsequent analysis. The raw signals at 5 kHz were downsampled to 100 Hz using an 8-pole bessel filter. Estimation of current jumps were carried out using a custom script with appropriate thresholds.

#### Estimation of dissociation constant (Kd) from nanopore conductance measurements

The analysis of nanopore data was generally following previously published protocols(*48*). The following assumptions were made for estimating the Kd of ligand binding to the CHD nanopore sensor. First, despite the presence of three distinct current states of the nanopore in the presence of ligand, only the lowest current level was considered to be indicative of ligand binding. Therefore current level jumps from the lowest state to either of the higher current states were treated identically and were considered to be indicative of unbinding. Second, ligand binding was assumed to stabilize the binder domain in a closed conformation and likely prolong the lowest current state of the nanopore. However, since the lowest current state is also observed for the control condition with no ligand, it was assumed that the dwell time of this current state in the presence of ligand would be a combination of inherent stochasticity of the binder domain closing and potential ligand binding.

With the above assumptions, a hill equation model with hill coefficient as one was sufficient to capture the binder-domain transitions from an unbound to a ligand-bound state:

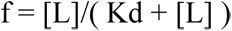

Here f denotes the fraction of the ligand bound state of the nanopore and [L] is the concentration of the ligand in solution which was fixed at 10 uM for this experiment.

Dwell time distribution for events resulting in the lowest current state of the nanopore were calculated for both cases (absence and presence of ligand) and were fitted to an exponential curve to estimate the mean dwell times (inverse of the exponential decay constants), Tc (nanopore only) and Tl (nanopore with ligand) respectively (Figure S17c). The mean dwell times were normalized by the duration of the recordings to represent the probabilities of the unbound and bound states. To prevent user bias in determining a cutoff for discarding very small dwell times resulting from noise or initial signal-filtering artifacts, a range of dwell time cutoffs from 50 ms to 400 ms were used. This resulted in a range of mean dwell times post exponential curve-fitting for each experimental condition. For each pair of Tc and Tl, the fraction of ligand-bound receptors was estimated as:

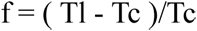

Corresponding Kd values were calculated with [L] = 10 uM. The range of calculated Kd is shown as a histogram in Figure S17d.

#### Engineering/Buttressing pseudocycle binders

The overall steps are shown in **Fig S18a**. The starting pseudocycle binder was first broken into individual repeat units based on their structural symmetry order and expanded to multiple perfect symmetric structures. We mutated the surface residues to apolar residues to help diffuse buttrased structures out. We also found use of ‘adjacent_matrices’ helps placement of buttressed structures. Then, we used symmetry diffusion to generate buttress around the individual symmetric structures(*10*). Symmetric diffusion was used because it is significantly faster and easier to converge to high quality protein scaffolds than free diffusion. We then take individual parts of the buttressed unit, mixed and matched them together to make buttressed binders. The binding interface residues were not designed while the rest of the residues were redesigned using proteinMPNN(*13*), scored with Rosetta, and folded using AF2(*14*). The 68 buttrased designs passed Rosetta, AF2 filters were ordered using Twist gene fragments. The designs were first transformed into EBY1 with pETCON3 vector for yeast surface display test to identify initial binders (see method ‘***Yeast surface display experiment for enrichment of the potential binders***‘). The designs with potential binding signal on yeast surface display were purified following protocol ‘***Expression and purification of selected proteins***’, and the binding affinity was determined using FP following protocol ‘***Fluorescence polarization studies***’.

### CID design and testing

#### Computational design of CID

The scripts for CID design were uploaded to github: https://github.com/iamlongtran/pseudocycle_paper (will change from private to public upon BiorXiv release).

The design of CID starts with verified buttressed binders, which were splitted into two halves (A and B). The main objective is to keep the ligand binding in presence of the SM ligand, while preventing the A-B interactions without the ligand. We first used ligandMPNN(*16*) for sequence design. Specifically, we kept the verified ligand-protein interface, and used tied sequence design functions, which allows simultaneous sequence design on A-ligand complex, B-ligand complex, and A-ligand-B complex, and used tied weight to reward A-ligand-B complex design and weak the rest conformation design.

Once we have designed CIDs, we predicted the A-B dimer formation using superfold, an in-house AF2 parser (https://github.com/rdkibler/superfold) to select designs which were predicted not to form hetero/homo-dimers but form stable individual monomeric proteins.

For A-ligand-B complexes, we use RosettaFold All Atom (RFAA)(*9*) to predict the complex structure. We selected designs with Predicted Aligned Error (PAE) lower than 5 and predicted model with low Cɑ-r.m.s.d to designs, and ordered as gene fragments from Twist Bioscience.

#### Chromatography test for identification of CID pairs

The expression and purification protocol of all proteins are closely followed to the section ‘***Expression and purification of selected proteins*’**, with a few minor adjustments. For the volume of autoinduction TB media, for CID screening, designs were grown in 5 mL TB autoinduction media in Falcon™ Round-Bottom Polypropylene Test Tubes with Cap in replicates. After growing for 6 hours at 37 ℃ followed by 24 hours at 18 ℃ with shaking at 225 rpm, cultures of each design were transferred to a separate well from a 24-well plate. Design cultures were harvested using centrifugation at 4000 g for 15 minutes, then resuspended in a 10 mL lysis buffer in the 24-well plate. The suspended cells were sonicated to lyse for 7.5 minutes, using 10 second pulse, 10 second pause, at 65% amplitude. The lyzed mixtures in 24-well collection plates were then centrifuged at 2000 g for 20 minutes to separate cell debris and lysate. The soluble fraction collected post centrifugation were bound to 1 mL of Ni-NTA resin in a 24-well filter plate, followed by wash and elution facilitated by a vacuum manifesto. The individual eluted proteins were first purified by SEC, and the CID pairs were analyzed on an Agilent 1260 Infinity I HPLC system with DAD using an analytical SEC column, superdex 75 increase 3.2/300 GL column. An 4-minute isocratic run was performed for each CID part and CID combinations to identify CID pairs.

#### Mass photometry test for identification of CID pairs

The mass photometry assays were performed following previously published protocol(*49*). All mass photometry measurements were carried out in an instrument TwoMP (Refeyn) Mass photometer. The same buffer (20 mM Tris 100 mM NaCl, pH 8) was used for all mass photometry experiments. Due to the small size of the CID monomers (A is 9.8 kDa, B is 18.9 kDa), we fused N-terminus gfp to each monomer to increase their molecular weight so that we can observe them under mass photometry. CID pairs were prepared as 10 nM protein gfp-A, gfp-B, with and without 10 μM ligand, and were used for all measurements after 1 hour incubation at r.t. A 24-well gasket was placed on a clean glass slide as sample holders. A 5 μL buffer was first added to one well of this gasket and used to bring the camera into focus after orienting the laser to the center of the sample well. A 5 μL sample prepared as stated-above was added to this droplet and 1-minute videos were collected with a large field of view in AcquireMP. Ratiometric contrast values for individual particles were measured and processed into mass distributions with DiscoverMP based on calibration curves generated with 20 nM β-amylase (consists of monomer 56 kDa, dimer 112 kDa, and tetramer 224 kDa). The dimerized peak of gfp-A and gfp-B in presence of ligand was clearly observed with a clear right shift compared to the monomer-only peaks and a good mass fitting based on the standard curve. The monomer peak of gfp-A, gfp-B was observed, clearly smaller than the dimerized peak, but cannot be fitted to report mass value due to their small size (gfp-A: 37 kDa, gfp-B: 46 kDa).

#### Crystallization condition and analysis

Crystallization experiment for the designed protein was conducted using the sitting drop vapor diffusion method. Crystallization trials were set up in 200 nL drops using the 96-well plate format at 20°C. Crystallization plates were set up using a Mosquito LCP from SPT Labtech, then imaged using UVEX microscopes from JAN Scientific. Diffraction quality crystals formed in 0.1 M Tris/Biocine pH 8.5, 25% v/v 2-methyl-2,4-pentanediol (MPD); 25% PEG1000; 25% w/v PEG 3350 and 0.093 MSodium fluoride; 0.3 M Sodium bromide; 0.3 M Sodium iodide for CHD_r1. CHD_r1_buttress crystal formed in 0.2 M Sodium chloride 20% (w/v) PEG 3350.

Diffraction data were collected at Advanced Photon Source on Beamline NECAT 24IDC for CHD_r1 and CHD_r1_buttress. X-ray intensities and data reduction were evaluated and integrated using XDS(*50*) and merged/scaled using Pointless/Aimless in the CCP4 program suite(*51*). Structure determination and refinement starting phases were obtained by molecular replacement using Phaser(*52*) using the designed model structure. Following molecular replacement, the models were improved using phenix.autobuild(*53*). Structures were refined in Phenix(*53*). Model building was performed using COOT(*54*). The final model was evaluated using MolProbity(*55*). Data collection and refinement statistics are recorded in the Supplementary **Table S3**. Data deposition, atomic coordinates, and structure factors reported for the protein in this paper have been deposited in the Protein Data Bank (PDB), http://www.rcsb.org/ with accession code 8VEI and 8VEJ.

## Reference

1. A. M. Gao, J. Skolnick, The distribution of ligand-binding pockets around protein-protein interfaces suggests a general mechanism for pocket formation. Proc. Natl. Acad. Sci. U. S. 109, 3784–3789 (2012).

2. J. Takeuchi, K. Fukui, Y. Seto, Y. Takaoka, M. Okamoto, Ligand–receptor interactions in plant hormone signaling. Plant J. 105, 290–306 (2021).

3. J. Dou, A. A. Vorobieva, W. Sheffler, L. A. Doyle, H. Park, M. J. Bick, B. Mao, G. W. Foight, M. Y. Lee, L. A. Gagnon, L. Carter, B. Sankaran, S. Ovchinnikov, E. Marcos, P.-S. Huang, J. C. Vaughan, B. L. Stoddard, D. Baker, De novo design of a fluorescence-activating β-barrel. Nature 561, 485–491 (2018).

4. J. Dou, L. Doyle, P. Greisen Jr., A. Schena, H. Park, K. Johnsson, B. L. Stoddard, D. Baker, Sampling and energy evaluation challenges in ligand binding protein design: Computational Protein Design. Protein Sci. 26, 2426–2437 (2017).

5. M. J. Bick, P. J. Greisen, K. J. Morey, M. S. Antunes, D. La, B. Sankaran, L. Reymond, K. Johnsson, J. I. Medford, D. Baker, Computational design of environmental sensors for the potent opioid fentanyl. eLife 6, e28909 (2017).

6. N. F. Polizzi, W. F. DeGrado, A defined structural unit enables de novo design of small-molecule-binding proteins. Science 369, 1227–1233 (2020).

7. C. E. Tinberg, S. D. Khare, J. Dou, L. Doyle, J. W. Nelson, A. Schena, W. Jankowski, C. G. Kalodimos, K. Johnsson, B. L. Stoddard, D. Baker, Computational design of ligand-binding proteins with high affinity and selectivity. Nature 501, 212–216 (2013).

8. G. R. Lee, S. J. Pellock, C. Norn, D. Tischer, J. Dauparas, I. Anishchenko, J. A. M. Mercer, A. Kang, A. Bera, H. Nguyen, I. Goreshnik, D. Vafeados, N. Roullier, H. L. Han, B. Coventry, H. K. Haddox, D. R. Liu, A. H.-W. Yeh, D. Baker, Small-molecule binding and sensing with a designed protein family. doi: 10.1101/2023.11.01.565201 (2023).

9. R. Krishna, J. Wang, W. Ahern, P. Sturmfels, P. Venkatesh, I. Kalvet, G. R. Lee, F. S. Morey-Burrows, I. Anishchenko, I. R. Humphreys, R. McHugh, D. Vafeados, X. Li, G. A. Sutherland, A. Hitchcock, C. N. Hunter, M. Baek, F. DiMaio, D. Baker, Generalized Biomolecular Modeling and Design with RoseTTAFold All-Atom. doi: 10.1101/2023.10.09.561603 (2023).

10. J. L. Watson, D. Juergens, N. R. Bennett, B. L. Trippe, J. Yim, H. E. Eisenach, W. Ahern, A. J. Borst, R. J. Ragotte, L. F. Milles, B. I. M. Wicky, N. Hanikel, S. J. Pellock, A. Courbet, W. Sheffler, J. Wang, P. Venkatesh, I. Sappington, S. V. Torres, A. Lauko, V. De Bortoli, E. Mathieu, R. Barzilay, T. S. Jaakkola, F. DiMaio, M. Baek, D. Baker, “Broadly applicable and accurate protein design by integrating structure prediction networks and diffusion generative models” (preprint, Biochemistry, 2022); bioRxiv.

11. L. An, D. R. Hicks, D. Zorine, J. Dauparas, B. I. M. Wicky, L. F. Milles, A. Courbet, A. K. Bera, H. Nguyen, A. Kang, L. Carter, D. Baker, Hallucination of closed repeat proteins containing central pockets. Nat. Struct. Mol. Biol., doi: 10.1038/s41594-023-01112-6 (2023).

12. A. Leaver-Fay, M. Tyka, S. M. Lewis, O. F. Lange, J. Thompson, R. Jacak, K. W. Kaufman, P. D. Renfrew, C. A. Smith, W. Sheffler, I. W. Davis, S. Cooper, A. Treuille, D. J. Mandell, F. Richter, Y.-E. A. Ban, S. J. Fleishman, J. E. Corn, D. E. Kim, S. Lyskov, M. Berrondo, S. Mentzer, Z. Popović, J. J. Havranek, J. Karanicolas, R. Das, J. Meiler, T. Kortemme, J. J. Gray, B. Kuhlman, D. Baker, P. Bradley, “Rosetta3” in Methods in Enzymology (Elsevier, 2011; https://linkinghub.elsevier.com/retrieve/pii/B9780123812704000196) vol. 487, pp. 545–574.

13. J. Dauparas, I. Anishchenko, N. Bennett, H. Bai, R. J. Ragotte, L. F. Milles, B. I. M. Wicky, Courbet, R. J. de Haas, N. Bethel, P. J. Y. Leung, T. F. Huddy, S. Pellock, D. Tischer, F. Chan, B. Koepnick, H. Nguyen, A. Kang, B. Sankaran, A. K. Bera, N. P. King, D. Baker, Robust deep learning-based protein sequence design using ProteinMPNN. Science 378, 49–56 (2022).

14. J. Jumper, R. Evans, A. Pritzel, T. Green, M. Figurnov, O. Ronneberger, K. Tunyasuvunakool, R. Bates, A. Žídek, A. Potapenko, A. Bridgland, C. Meyer, S. A. A. Kohl, A. J. Ballard, A. Cowie, B. Romera-Paredes, S. Nikolov, R. Jain, J. Adler, T. Back, S. Petersen, D. Reiman, E. Clancy, M. Zielinski, M. Steinegger, M. Pacholska, T. Berghammer, S. Bodenstein, D. Silver, O. Vinyals, A. W. Senior, K. Kavukcuoglu, P. Kohli, D. Hassabis, Highly accurate protein structure prediction with AlphaFold. Nature 596, 583–589 (2021).

15. M. Baek, F. DiMaio, I. Anishchenko, J. Dauparas, S. Ovchinnikov, G. R. Lee, J. Wang, Q. Cong, L. N. Kinch, R. D. Schaeffer, C. Millán, H. Park, C. Adams, C. R. Glassman, A. DeGiovanni, J. H. Pereira, A. V. Rodrigues, A. A. van Dijk, A. C. Ebrecht, D. J. Opperman, T. Sagmeister, C. Buhlheller, T. Pavkov-Keller, M. K. Rathinaswamy, U. Dalwadi, C. K. Yip, J. E. Burke, K. C. Garcia, N. V. Grishin, P. D. Adams, R. J. Read, D. Baker, Accurate prediction of protein structures and interactions using a three-track neural network. Science 373, 871–876 (2021).

16. J. Duparas, G. R. Lee, R. Pecoraro, L. An, I. Anishchenko, C. Glasscock, D. Baker, Atomic context-conditioned protein sequence design using LigandMPNN. Biorxiv.

17. X. Zhao, Z. Liu, F. Sun, L. Yao, G. Yang, K. Wang, Bile Acid Detection Techniques and Bile Acid-Related Diseases. Front. Physiol. 13, 826740 (2022).

18. A. Shadeed, L. Kattach, S. Sam, K. Flora, Z. Farah, Examining the safety of relaxed drug monitoring for methotrexate in response to the COVID-19 pandemic. Rheumatol. Adv. Pract. 6, rkac100 (2022).

19. C. A. Spencer, Assay of Thyroid Hormones and Related Substances (Endotext [Internet], South Dartmouth (MA), 2017; https://www.ncbi.nlm.nih.gov/books/NBK279113/).

20. K. J. Welsh, S. J. Soldin, DIAGNOSIS OF ENDOCRINE DISEASE: How reliable are free thyroid and total T3 hormone assays? Eur. J. Endocrinol. 175, R255–R263 (2016).

21. N. K. Tripathy, S. K. Mishra, G. Nathan, S. Srivastava, A. Gupta, R. Lingaiah, A Rapid Method for Determination of Serum Methotrexate Using Ultra-High-Performance Liquid Chromatography-Tandem Mass Spectrometry and Its Application in Therapeutic Drug Monitoring. J. Lab. Physicians 15, 344–353 (2023).

22. P. Salveson, De novo design f macrocycles.

23. G. Landrum, P. Tosco, B. Kelley, Ric, D. Cosgrove, Sriniker, Gedeck, R. Vianello, NadineSchneider, E. Kawashima, D. N. G. Jones, A. Dalke, B. Cole, M. Swain, S. Turk, AlexanderSavelyev, A. Vaucher, M. Wójcikowski, Ichiru Take, D. Probst, K. Ujihara, V. F. Scalfani, G. Godin, J. Lehtivarjo, R. Walker, A. Pahl, Francois Berenger, Jasondbiggs, Strets123, rdkit/rdkit: 2023_03_3 (Q1 2023) Release, version Release_2023_03_3, Zenodo (2023); 10.5281/ZENODO.591637.

24. Sagadip, De novo nanopore. Biorxiv.

25. J. Wang, S. Lisanza, D. Juergens, D. Tischer, J. L. Watson, K. M. Castro, R. Ragotte, A. Saragovi, L. F. Milles, M. Baek, I. Anishchenko, W. Yang, D. R. Hicks, M. Expòsit, T. Schlichthaerle, J.-H. Chun, J. Dauparas, N. Bennett, B. I. M. Wicky, A. Muenks, F. DiMaio, B. Correia, S. Ovchinnikov, D. Baker, Scaffolding protein functional sites using deep learning. Science 377, 387–394 (2022).

26. M. Ahmad, J.-H. Ha, L. A. Mayse, M. F. Presti, A. J. Wolfe, K. J. Moody, S. N. Loh, L. Movileanu, A generalizable nanopore sensor for highly specific protein detection at single-molecule precision. Nat. Commun. 14, 1374 (2023).

27. M. A. Fahie, M. Chen, Electrostatic Interactions between OmpG Nanopore and Analyte Protein Surface Can Distinguish between Glycosylated Isoforms. J. Phys. Chem. B 119, 10198–10206 (2015).

28. J. C. Foster, B. Pham, R. Pham, M. Kim, M. D. Moore, M. Chen, An Engineered OmpG Nanopore with Displayed Peptide Motifs for Single-Molecule Multiplex Protein Detection. Angew. Chem. Int. Ed Engl. 62, e202214566 (2023).

29. B. L. Greene, G. Kang, C. Cui, M. Bennati, D. G. Nocera, C. L. Drennan, J. Stubbe, Ribonucleotide Reductases: Structure, Chemistry, and Metabolism Suggest New Therapeutic Targets. Annu. Rev. Biochem. 89, 45–75 (2020).

30. M. Rappas, H. Niwa, X. Zhang, Mechanisms of ATPases - A Multi-Disciplinary Approach. Curr. Protein Pept. Sci. 5, 89–105 (2004).

31. L. A. Banaszynski, C. W. Liu, T. J. Wandless, Characterization of the FKBP·Rapamycin·FRB Ternary Complex. J. Am. Chem. Soc. 127, 4715–4721 (2005).

32. G. W. Foight, Z. Wang, C. T. Wei, P. Jr Greisen, K. M. Warner, D. Cunningham-Bryant, K. Park, T. J. Brunette, W. Sheffler, D. Baker, D. J. Maly, Multi-input chemical control of protein dimerization for programming graded cellular responses. Nat. Biotechnol. 37, 1209–1216 (2019).

33. B. Z. Stanton, E. J. Chory, G. R. Crabtree, Chemically induced proximity in biology and medicine. Science 359, eaao5902 (2018).

34. J. Shen, Y. Gu, L. Ke, Q. Zhang, Y. Cao, Y. Lin, Z. Wu, C. Wu, Y. Mu, Y.-L. Wu, C. Ren, H. Zeng, Cholesterol-stabilized membrane-active nanopores with anticancer activities. Nat. Commun. 13, 5985 (2022).

35. L. Holm, A. Laiho, P. Törönen, M. Salgado, DALI shines a light on remote homologs: One hundred discoveries. Protein Sci. Publ. Protein Soc. 32, e4519 (2023).

36. S. Kim, J. Chen, T. Cheng, A. Gindulyte, J. He, S. He, Q. Li, B. A. Shoemaker, P. A. Thiessen, B. Yu, L. Zaslavsky, J. Zhang, E. E. Bolton, PubChem 2023 update. Nucleic Acids Res. 51, D1373–D1380 (2023).

37. N. M. O’Boyle, M. Banck, C. A. James, C. Morley, T. Vandermeersch, G. R. Hutchison, Open Babel: An open chemical toolbox. J. Cheminformatics 3, 33 (2011).

38. W. Humphrey, A. Dalke, K. Schulten, VMD: Visual molecular dynamics. J. Mol. Graph. 14, 33–38 (1996).

39. H. Park, G. Zhou, M. Baek, D. Baker, F. DiMaio, Force Field Optimization Guided by Small Molecule Crystal Lattice Data Enables Consistent Sub-Angstrom Protein-Ligand Docking. J. Chem. Theory Comput. 17, 2000–2010 (2021).

40. X. Qi, F. Loiseau, W. L. Chan, Y. Yan, Z. Wei, L.-G. Milroy, R. M. Myers, S. V. Ley, R. J. Read, R. W. Carrell, A. Zhou, Allosteric Modulation of Hormone Release from Thyroxine and Corticosteroid-binding Globulins. J. Biol. Chem. 286, 16163–16173 (2011).

41. D. M. Hoover, DNAWorks: an automated method for designing oligonucleotides for PCR-based gene synthesis. Nucleic Acids Res. 30, 43e–443 (2002).

42. G. Chao, W. L. Lau, B. J. Hackel, S. L. Sazinsky, S. M. Lippow, K. D. Wittrup, Isolating and engineering human antibodies using yeast surface display. Nat. Protoc. 1, 755–768 (2006).

43. L. Cao, B. Coventry, I. Goreshnik, B. Huang, W. Sheffler, J. S. Park, K. M. Jude, I. Marković, R. U. Kadam, K. H. G. Verschueren, K. Verstraete, S. T. R. Walsh, N. Bennett, A. Phal, A. Yang, L. Kozodoy, M. DeWitt, L. Picton, L. Miller, E.-M. Strauch, N. D. DeBouver, A. Pires, A. K. Bera, S. Halabiya, B. Hammerson, W. Yang, S. Bernard, L. Stewart, I. A. Wilson, H. Ruohola-Baker, J. Schlessinger, S. Lee, S. N. Savvides, K. C. Garcia, D. Baker, Design of protein-binding proteins from the target structure alone. Nature 605, 551–560 (2022).

44. B. Dang, M. Mravic, H. Hu, N. Schmidt, B. Mensa, W. F. DeGrado, SNAC-tag for sequence-specific chemical protein cleavage. Nat. Methods 16, 319–322 (2019).

45. L. An, D. P. Cogan, C. D. Navo, G. Jiménez-Osés, S. K. Nair, W. A. van der Donk, Substrate-assisted enzymatic formation of lysinoalanine in duramycin. Nat. Chem. Biol. 14, 928–933 (2018).

46. A. A. Vorobieva, De novo design of nanopores. Biorxiv.

47. A. A. Vorobieva, P. White, B. Liang, J. E. Horne, A. K. Bera, C. M. Chow, S. Gerben, S. Marx, A. Kang, A. Q. Stiving, S. R. Harvey, D. C. Marx, G. N. Khan, K. G. Fleming, V. H. Wysocki, D. J. Brockwell, L. K. Tamm, S. E. Radford, D. Baker, De novo design of transmembrane β barrels. Science 371, eabc8182 (2021).

48. N. Varongchayakul, J. Song, A. Meller, M. W. Grinstaff, Single-molecule protein sensing in a nanopore: a tutorial. Chem. Soc. Rev. 47, 8512–8524 (2018).

49. A. Pillai, A. Idris, A. Philomin, C. Weidle, R. Skotheim, P. J. Y. Leung, A. Broerman, C. Demakis, A. J. Borst, F. Praetorius, D. Baker, “De novo design of allosterically switchable protein assemblies” (preprint, Biochemistry, 2023); 10.1101/2023.11.01.565167.

50. W. Kabsch, XDS. Acta Crystallogr. D Biol. Crystallogr. 66, 125–132 (2010).

51. M. D. Winn, C. C. Ballard, K. D. Cowtan, E. J. Dodson, P. Emsley, P. R. Evans, R. M. Keegan, E. B. Krissinel, A. G. W. Leslie, A. McCoy, S. J. McNicholas, G. N. Murshudov, N. S. Pannu, E. A. Potterton, H. R. Powell, R. J. Read, A. Vagin, K. S. Wilson, Overview of the *CCP* 4 suite and current developments. Acta Crystallogr. D Biol. Crystallogr. 67, 235–242 (2011).

52. A. J. McCoy, R. W. Grosse-Kunstleve, P. D. Adams, M. D. Winn, L. C. Storoni, R. J. Read, Phaser crystallographic software. J. Appl. Crystallogr. 40, 658–674 (2007).

53. P. D. Adams, P. V. Afonine, G. Bunkóczi, V. B. Chen, I. W. Davis, N. Echols, J. J. Headd, L.-W. Hung, G. J. Kapral, R. W. Grosse-Kunstleve, A. J. McCoy, N. W. Moriarty, R. Oeffner, R. J. Read, D. C. Richardson, J. S. Richardson, T. C. Terwilliger, P. H. Zwart, *PHENIX*: a comprehensive Python-based system for macromolecular structure solution. Acta Crystallogr. D Biol. Crystallogr. 66, 213–221 (2010).

54. P. Emsley, K. Cowtan, Coot: model-building tools for molecular graphics. Acta Crystallogr. D Biol. Crystallogr. 60, 2126–2132 (2004).

55. C. J. Williams, J. J. Headd, N. W. Moriarty, M. G. Prisant, L. L. Videau, L. N. Deis, V. Verma, D. A. Keedy, B. J. Hintze, V. B. Chen, S. Jain, S. M. Lewis, W. B. Arendall, J. Snoeyink, P. D. Adams, S. C. Lovell, J. S. Richardson, D. C. Richardson, MolProbity: More and better reference data for improved all-atom structure validation: PROTEIN SCIENCE.ORG. *Protein Sci.* 27, 293–315 (2018).

